# Xolography for Biomedical Applications: Dual-color Light-sheet Printing of Hydrogels with Local Control over Shape and Stiffness

**DOI:** 10.1101/2024.12.21.629893

**Authors:** Lena Stoecker, Gerardo Cedillo-Servin, Niklas F. König, Freek V. de Graaf, Marcela García Jiménez, Sandra Hofmann, Keita Ito, Annelieke S. Wentzel, Miguel Castilho

## Abstract

Current challenges in tissue engineering include creation of extracellular environments that support and interact with cells using biochemical, mechanical, and structural cues. Spatial control over these cues is currently limited due to a lack of suitable fabrication techniques. This study introduces Xolography, an emerging dual-color light-sheet volumetric printing technology, to achieve control over structural and mechanical features for hydrogel-based photoresins at micro-to macroscale while printing within minutes. We propose a water-soluble photoswitch photoinitiator system and are the first to demonstrate Xolography with a library of naturally-derived, synthetic, and thermoresponsive hydrogels. Centimeter-scale, three-dimensional constructs with positive features of 20 µm and negative features of ∼ 100 µm are fabricated with control over mechanical properties (compressive moduli 0.2 kPa – 6.5 MPa). Notably, switching from binary to grayscaled light projection enables spatial control over stiffness (0.2 – 16 kPa). As a proof of concept, grayscaled Xolography is leveraged with thermoresponsive hydrogels to introduce reversible anisotropic shape changes beyond isometric shrinkage. We finally demonstrate Xolography of viable cell aggregates, laying the foundation for cell-laden printing of dynamic, cell-instructive environments with tunable structural and mechanical cues in a fast one-step process. Overall, these innovations unlock unique possibilities of Xolography across multiple biomedical applications.

## 1. Introduction

Tissue engineering aims to design extracellular environments that support and instruct cells and thus relies on materials that provide diverse biochemical, mechanical, and structural cues. Hydrogels are the most widely used material class for this purpose because of their biocompatibility, biodegradability, and capability to retain large amounts of water. Furthermore, biochemical and mechanical properties of hydrogels can generally be targeted towards specific tissues by tuning the base polymer composition, water content, as well as type and degree of crosslinking (DoC). While many hydrogels are able to provide an environment that cells can remodel, traditional hydrogel fabrication modalities often lack precise control over structural aspects (e.g. porosity, geometrical shape, anisotropy), mechanical heterogeneity (e.g. local elasticity, local viscoelasticity), and dynamic behavior (e.g. degradation rate, volumetric swelling, actuation), which are crucial to instruct cell fate and guide tissue development over space and time.^[1–4]^

Additive manufacturing (AM) is increasingly utilized to address this limitation. AM offers improved design freedom compared to formative and subtractive manufacturing and is therefore suited to assign complex geometrical shapes to hydrogels. Light-based additive manufacturing, classified as vat photopolymerization, is of increasing interest for tissue engineering due to its superior feature resolution compared to extrusion and jetting processes and its ability for spatiotemporal control.^[5]^ Two-photon polymerization (TPP), for instance, has enabled feature resolution on sub-cellular level to create patterns that direct cell morphology.^[6, 7]^ However, low building rates limit the use of TPP for millimeter-and centimeter-scale constructs. Other vat photopolymerization technologies based on multi-directional or tomographic projections allow for higher building rates which facilitate production of tissue-level constructs.^[8–10]^ They lack nanoscale feature resolution but offer resolution on cellular-level (25–100 µm). While multi-directional AM has not yet been applied for cell-laden printing, tomographic AM has recently shown promising potential for fabricating tissue-like structures.^[11, 12]^ This process relies on the overlay of segmented images in a rotating cuvette which cumulatively reach sufficient energy to activate photopolymerization. Heterogeneity in the energy dosage can occur due to the mathematically determined segmentation, especially for sharp edges or hollow geometries perpendicular to the axis of rotation. This causes crosslinking differences throughout the print, so *in situ* control of the DoC and subsequent mechanical properties are yet to be addressed. A stiffness gradient from 66 MPa to 1.7 GPa was achieved within a distance of 300 µm for non-biocompatible photoresins but required addition of a supplementary photoresin.^[13]^

In turn, Filamented Light (FLight) biofabrication is an AM process that demonstrates that mechanical heterogeneity inside a hydrogel is capable of instructing cell alignment by generating crosslinked microfiber regions within an initially homogeneous hydrogel precursor.^[14]^ Nevertheless, FLight lacks precise control over the printed geometry along the third dimension. Currently available AM technologies thus face limitations in simultaneously creating (i) high feature resolution at the cellular or sub-cellular level (nm–µm) in (ii) short processing times (<10 min) with (iii) spatially resolved mechanical properties (elasticity), which are key for guiding cell behavior.

With these challenges in mind, the introduction of light-sheet volumetric printing and its first-ever representative Xolography was a promising development which could improve upon spatial control of structural and mechanical properties.^[15–17]^ Xolography utilizes photoswitch-chemistry and a cartesian print-setup, which in turn is based on the intersection of a UV light sheet with wavelength λ_1_ = 375 nm and a visible light projection λ_2_ (Figure 1). A dual-color photoinitiator (DCPI) is transitioned from its dormant isomer characterized by a spiropyran moiety (A) into a metastable active isomer characterized by a merocyanine moiety (B) by UV excitation (λ_1_). With this active form, orthogonally applied visible light (λ_2_) initiates radical formation and local photopolymerization (C). Three-dimensional objects are generated by moving the material with print speed υ through the UV light sheet at a specific intensity *Ι* while adapting the orthogonal light projection’s focus accordingly. Xolography has been demonstrated to reach feature resolution of 10 μm and object volumes of up to 2 cm³ (height = 2 cm, width = 1 cm) within processing times of one minute for synthetic acrylate resins.^[18]^ This concept has been employed for continuous manufacturing, fabrication of green-part ceramics, and multimaterial printing by a two-step overprinting approach and the incorporation of reactive species.^[19, 20]^ However, suitability of Xolography for (cell-laden) hydrogel printing has not been shown yet.

**Figure 1.**
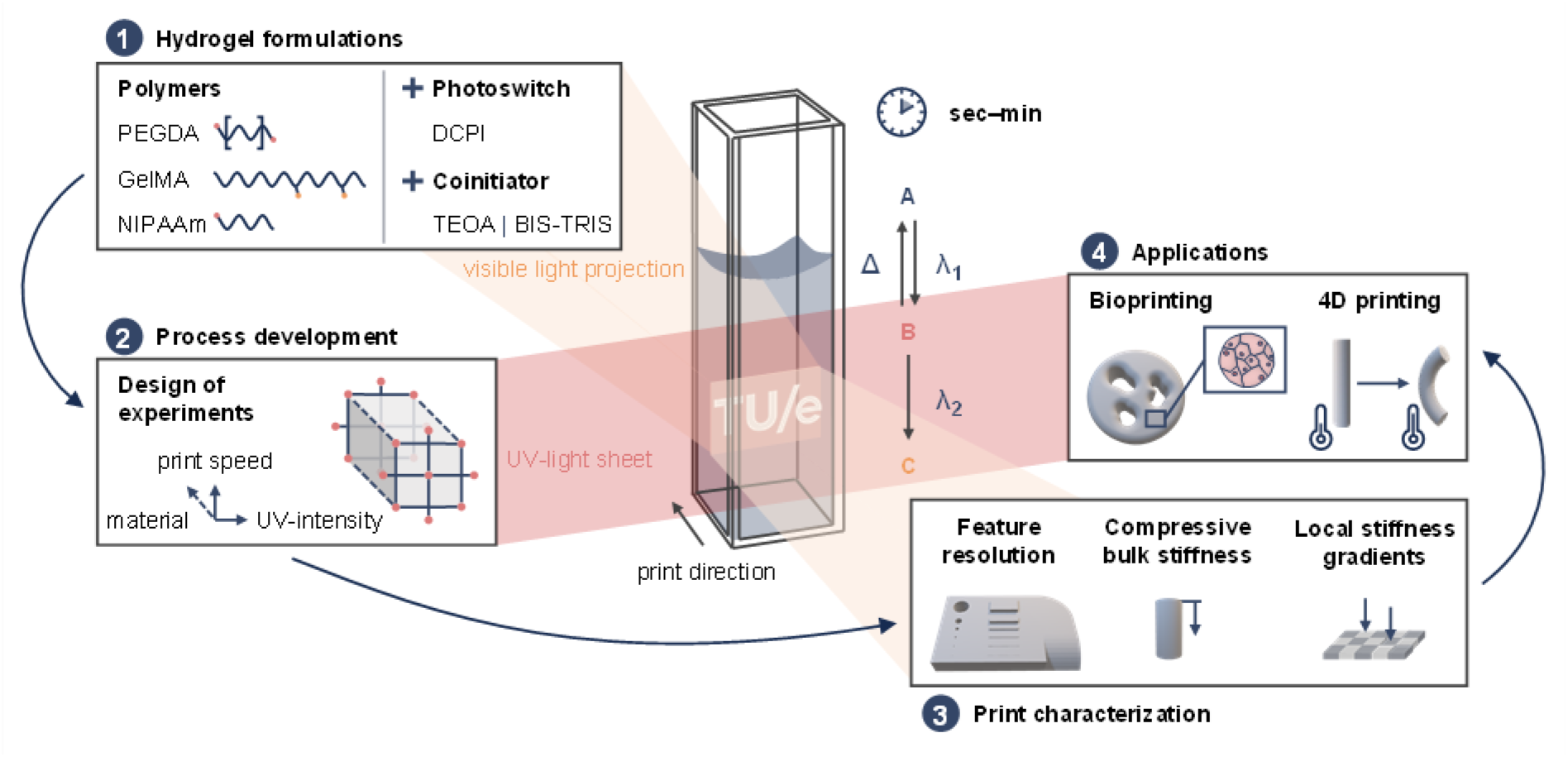
Schematic principle of local photopolymerization initiated by orthogonally applied light beams in Xolography (A = spiropyran, B = merocyanine, C = radical) and the steps undertaken to advance the technology for hydrogel manufacturing: (1) formulation of printable hydrogel systems, (2) process optimization, (3) characterization of printed specimens, and (4) demonstration of tissue engineering applications: bioprinting and 4D hydrogel printing.

We herein present the advancement of Xolography for fast manufacturing of hydrogels with microscale structural features and spatially resolved mechanical properties (Figure 1). We first design suitable photoresins based on a hydrophilic photoinitiator system and two hydrogels, gelatin methacryloyl (GelMA) and polyethylene glycol diacrylate (PEGDA), which are widely used in biomedical applications. We thoroughly investigate the printing process by design of experiments (DoE) and adjustment of hydrogel compositions, printing speed, and UV dosage.^[21]^ Furthermore, we showcase an additional level of spatial control by locally regulating the energy input for photopolymerization. By employing a grayscaled instead of a binary projection, we control material stiffness locally as a result of varying DoC. We then demonstrate that spatial modulation of DoC also enables 4D printing of hydrogels with dynamic, reversible shape-changing behavior by exploiting localized shrinkage differences in thermoresponsive photoresins based on *N*-isopropyl acrylamide (NIPAAm). Finally, we investigate toxicity of material formulations and demonstrate compatibility of the Xolography process for printing in the presence of cells by tracking the viability and behavior of cell-aggregates in printed scaffolds.

## 2. Results and discussion

### 2.1 Preparation of cytocompatible Xolography printing by selection of adequate DCPI and coinitiator

The development of a hydrogel-based photoresin for Xolography requires both adaptation of the photoinitiator system consisting of DCPI and coinitiator as well as selection of photopolymerizable hydrogels. The previously introduced DCPI molecule was composed of a benzophenone photoinitiator motif fused with a photochrome (t-type) photoswitch, which overall lacked aqueous solubility.^[17]^ The DCPI was thus functionalized with a water-solubilizing group and the further substitution pattern was optimized for the desired photo-physical properties in aqueous environment. The molecular structure of the adapted and water-soluble DCPI is schematically depicted in Figure 2. A The modified DCPI in its dormant form (spiropyran) absorbed light with wavelength below 400 nm (Figure 2 B), similar to its water-insoluble version.^[17]^ UV irradiation at 375 nm induced photoisomerization from spiropyran to merocyanine, as indicated by the characteristic merocyanine visible absorbance band around 560 nm. This validated the use of visible light as an orthogonal stimulus for the DCPI to induce polymerization. The merocyanine form nevertheless showed absorbance at 375 nm which opens a reaction channel for undesired UV light sheet-induced photopolymerization at high UV dosage.

**Figure 2.**
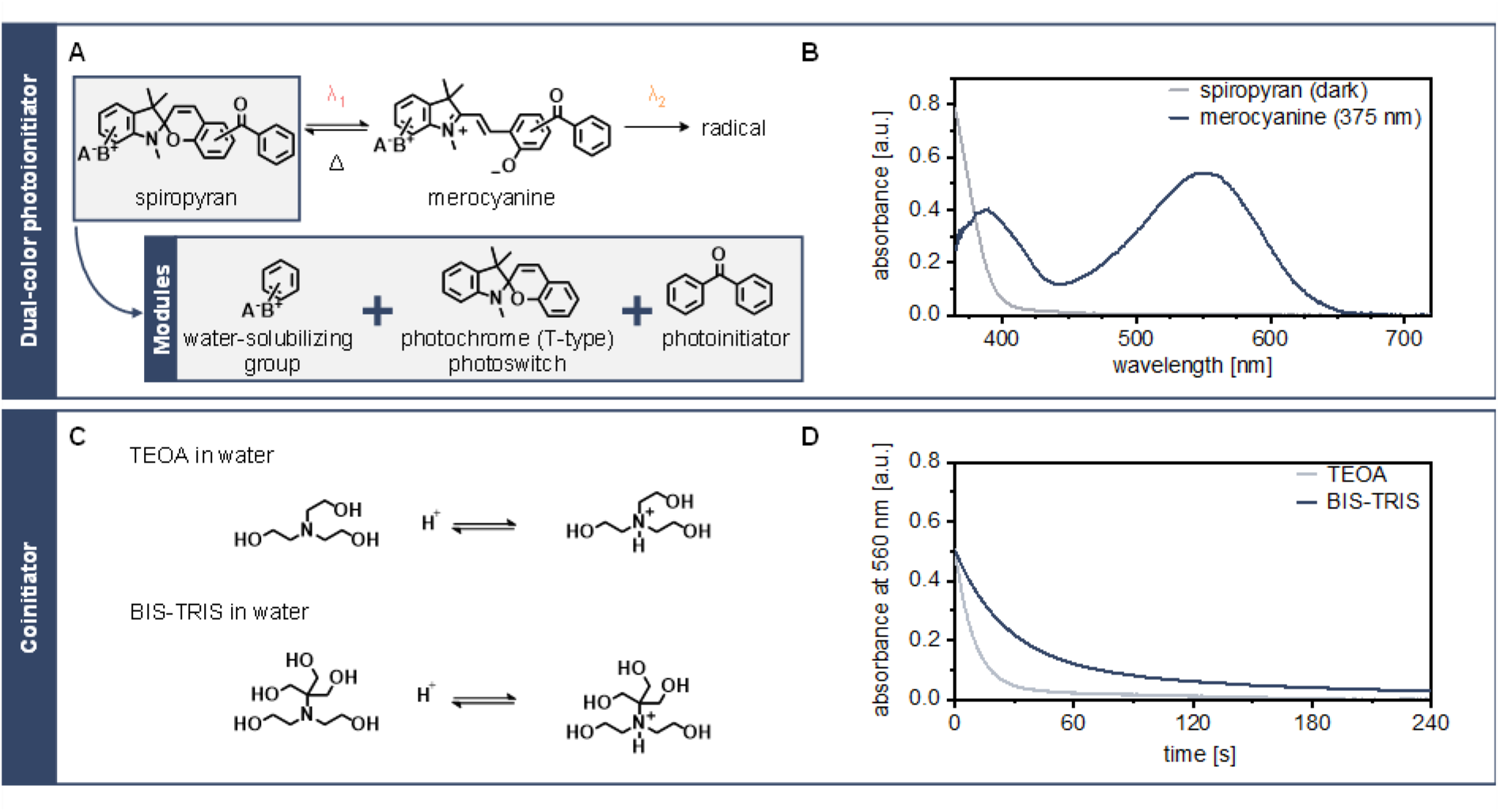
Characteristics of the adapted water-soluble photoinitiating system. (A) Modular chemical structure and reaction schematic of the hydrophilic DCPI. A^-^B^+^ denotes for an ionic group where either the cation or the anion can be covalently linked to the DCPI motif. (B) Absorbance spectrum of the hydrophilic DCPI in the dark and difference spectrum under irradiation at 375 nm, n = 1. (C) Chemical structures and their acid-base equilibria of employed coinitiators. (D) Thermal relaxation of the DCPI dependent on the employed coinitiator at a concentration of 800 mM, n = 1.

The DCPI belongs to the class of type II photoinitiators and requires a coinitiator, e.g. a tertiary amine: Upon excitation of the DCPI, an electron transfer from the amine nitrogen’s lone pair to the excited DCPI moiety is enabled, that can be followed by H-transfer, yielding an α-aminoalkyl radical to start a free radical polymerization.^[22]^ During the propagation reaction, covalent bonds are formed between reactive monomers resulting in long linear chains or crosslinked networks. Among the potential coinitiators, β-hydroxy-substituted tertiary alkylamines lead to highly nucleophilic and thus especially reactive α-amino radicals in the photoredox process allowing for an effective initiation.^[23]^

We successfully explored triethanolamine (TEOA) as coinitiator in synthetic hydrogels, as it has been successfully employed in Xolography of acrylate resins before.^[17]^ In the context of bioprinting however, TEOA is not an ideal choice as it is associated with cytotoxicity at concentrations above 0.05 mM.^[24]^ TEOA is moreover quite basic (pK_a_ = 7.76) and consequently predominantly protonated at the physiologically relevant pH of 7.4. The protonated tertiary amine is not an effective cointitator. For those reasons, we introduced the biocompatible Good’s buffer 2-bis(2-hydroxyethyl)amino-2-(hydroxymethyl)-1,3-propanediol (BIS-TRIS) as an alternative coinitiator in aqueous medium. (Figure 2 C) Unlike TEOA, BIS-TRIS is predominantly non-protonated and thus an active coinitiator at pH 7.4 due to its lower basicity (pK_a_ = 6.48).^[25]^ After its UV-induced formation, the activated DCPI merocyanine form shows a characteristic decay in the dark due to thermal isomerization back to the dormant spiropyran form. This thermal decay is pivotal to Xolography and influences the range of suitable printing speeds. Tracking of the characteristic visible light absorbance over time revealed that DCPI in TEOA and BIS-TRIS coinitiator solutions undergoes a biexponential decay with a short half-life of 6.6 s and longer half-life of 17.0 s, respectively (Figure 2 D). A fast decay is favorable for resolution in the print direction as it limits undesired crosslinking in areas just behind the moving UV light sheet.^[20]^ TEOA is therefore employed for photoresins targeting high resolution while BIS-TRIS is employed for photoresins that aim for cytocompatibility.

As hydrogel components, PEGDA is selected as a synthetic photopolymer which is commercially available with various average molecular numbers (M_n_). We expected fast reaction kinetics especially with low M_n_. Naturally-derived GelMA was included because PEGDA lacks any bioactive groups and GelMA allows for cell-attachment but suffers from lower reactivity, possibly due to high molecular weight (MW). We further employed a hybrid material formulation with the aim of combining the characteristics of both materials.

### 2.2 Xolography shows printability of synthetic and natural-derived hydrogel formulations

Crosslinking in light-based AM processes including Xolography is achieved if the energy input reaches a photoresin-specific activation threshold. Energy input is positively correlated to irradiation intensity and negatively correlated to printing speed. To optimize processability of hydrogel-based photoresins with Xolography, the effects of key processing parameters -printing speed and UV intensity - on feature resolution and mechanical stiffness were systematically investigated with DoE strategies. Initially, a full factorial screening of the parameter range was performed, followed by process analysis with response surface methodology and comparison to a previously established synthetic resin based on urethane dimethacrylate (UDMA).^[17]^

To investigate the capability of Xolography to manufacture hydrogel scaffolds with geometrical features on a cellular level, we introduced a test part containing negative and positive features in decreasing size and assessed their formation with different photoresins (Figure 3 E, Figure S1). Negative features in the range of 30 µm–1 mm represented channels and pores, similar in size to different biological structures like human vasculature or trabecular bone.^[26–28]^ Positive features provided insights into the minimum required height (20 µm–1 mm) of features to be printable (Figure S1). The design deviates from star-shaped geometries that are often used in literature in order to assess the still printable feature size along different directions which gives valuable insight for design processes.

**Figure 3.**
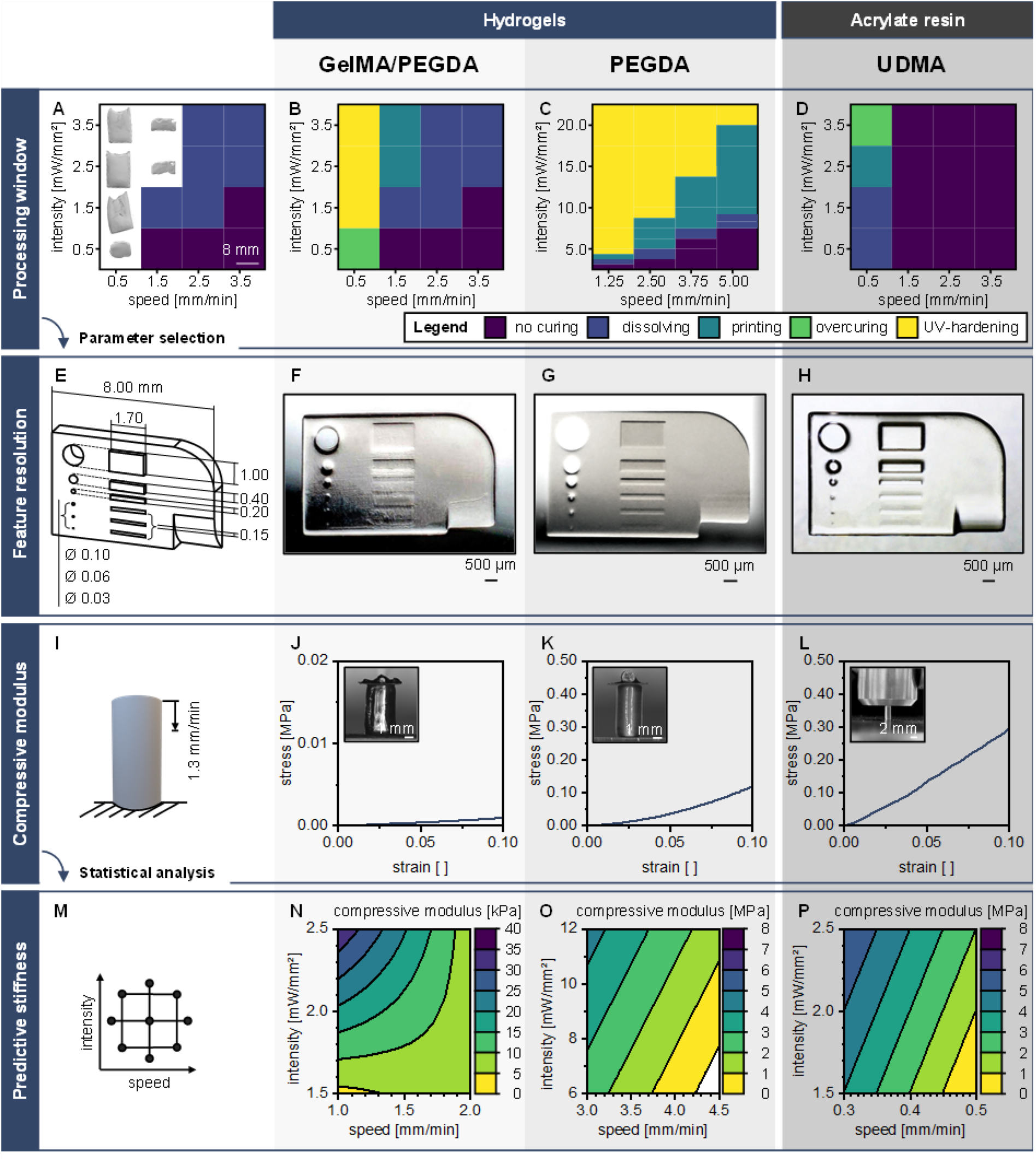
Systematic investigation of the processability of hydrogels with Xolography and the effect of printing speed and UV intensity on feature resolution and mechanical stiffness of printed parts in comparison to state-of-the-art acrylate resins. (A)–(D) Processing window of investigated material formulations based on prints of the geometry depicted in (E). (E)–(H) Resulting images from print optimization regarding resolution. (I)–(L) Compressive testing, and representative stress-strain curves accompanied by photographs of print optimization regarding resolution. (I)–(L) Compressive testing, and representative stress-strain curves accompanied by photographs of printed specimens under preload. (M)–(P) Prediction of achievable stiffness in the investigated parameter envelope. n = 1 samples (A–H), n = 3, two-way ANOVA (p < 0.05) (I–P), R^²^_adj_ = 63.1 % (N), R^²^_adj_ = 81.1 % (O), R^²^_adj_ = 76.2 % (P).

Full factorial parameter screening revealed suitable combinations of process parameters for the selected geometry. Figure 3 B–D shows the suitable parameter combinations for a GelMA/PEGDA, a PEGDA, and a UDMA photoresin (referred to as ‘printing’). The printing window is localized between parameter combinations that resulted insufficient crosslinking which led to dissolved parts during post-processing (referred to as ‘dissolving’), and combinations that caused crosslinking across the edges of the desired geometry due to overexposure (referred to as ‘overcuring’). All material formulations (GelMA/PEGDA, PEGDA, and UDMA) exhibited a region where energy input fell below the activation threshold due to a combination of low intensities and high printing speeds (referred to as ‘no curing’) and no crosslinking was observed. Additionally, hydrogel-based photoresins (GelMA/PEGDA, PEGDA) held a region where UV irradiation was sufficiently high to not only activate the photoswitch but also to initiate radical formation and photopolymerization. As a result, these areas prohibited precise formation of three-dimensional geometries and instead polymerized the entire cross-section of the cuvette (referred to as ‘UV hardening’). Figure 3 A visualizes the described phenomena for a GelMA/PEGDA photoresin. Sheet-like structures as seen for combinations of 1.5 mm min^-1^ printing speed and 1.5–3.5 mW mm^-^² UV intensity were typical for UV hardening. At 0.5 mm min^-1^ printing speed and 0.5 mW mm^-^² UV intensity, overcuring can be identified by an enlarged geometry in which the features of the intended geometry were barely distinguishable. Geometries with well-defined features were obtained at 1.5 mm min^-1^ printing speed and 2.5–3.5 mW mm^-^² UV intensity. Noticable differences between the CAD model and the printed features however demonstrated the need for the further described process optimization. Dissolving or no curing at all occurred for all combinations at 2.5–3.5 mm min^-1^ printing speed and 1.5 mm min^-1^ printing speed in combination with 0.5 and 1.5 mW mm^-^² UV intensity.

A pure GelMA photoresin was also tested but its printing window remained outside of the screened parameter range (data therefore not shown). Printability at speeds lower than 0.2 mm min^-1^ was likely due to a low fraction of primary amines (∼ 0.3 mmol g^-1^ of gelatin) which are the primary functionalization sites in gelatin in combination with a high MW (∼ 50–100 kDa) of the polymer.^[29, 30]^ This was in line with our observation that only GelMA photoresins with a degree of methacrylation (DoM) above 60 % could be printed, as well as with the previously reported dependency of crosslinking based on exposure time, intensity and DoM.^[29]^ The small processing window for GelMA resulted in limited potential for optimization for processing parameters and reduced printability of thin and delicate geometries such as the one in Figure 3 E. A hybrid hydrogel consisting of 10 % w/v GelMA and 10 % w/v PEGDA exhibited printability around a printing speed of 2.5 mm min^-1^ and intensities around 2.5–3.5 mW mm^-^² (Figure 3 B). The addition of PEGDA with a relatively small MW (M_n_ = 700) resulted in higher overall concentration of double bonds compared with GelMA alone.^[31]^ An evaluation of the crosslinking behavior for both GelMA and GelMA/PEGDA photoresins demonstrated that the crosslinking speed upon UV exposure of a hybrid hydrogel was 6 times faster than that of a pure GelMA-hydrogel (Figure 6 C).

UDMA showed a printing window only at the slowest examined printing speed of 0.5 mm min^-1^ and appeared more sensitive to changes in UV intensity, transitioning from dissolving to overcuring between 1.5–3.5 mW mm^-^² (Figure 3 D). The printing window of the PEGDA-photoresin was the largest of the materials investigated and adjustable to various printing speeds, with an increasing range of suitable intensities (9.0 –20 mW mm^-^²) at a high print speed (5 mm min^-1^).

Abovementioned energy input is frequently used to define printability in light-based AM. We found that the process variable energy input was not always sufficient to define printability of photoresins for Xolography. While a low printing speed of 0.5 mm min^-1^ coupled with low UV intensity of 0.5 mW mm^-^² led to overcured samples, the same energy input of 3 mJ mm^-^² as a combination of 1.5 mm min^-1^ and 1.5 mW mm^-^² resulted in dissolving samples (Table S1, Figure 3 B). This effect is likely enhanced by the two-step chemistry of the DCPI which requires both sufficient transitions of dormant to active isomers as well as sufficient radical formation within this excited DCPI.

We demonstrated that GelMA/PEGDA and PEGDA photoresins enabled printing of the test part (Figure 3 F, G). Notably, hydrogel-based photoresins demonstrated printability of positive features with heights as low as 20 µm and marks of all six channels, albeit only PEGDA-hydrogels showed pronounced edges and even represent angular approximation caused by the tessellation of the print file on the radius in x-z plane (Figure S1 A, Figure S2, Figure S3, Video S1). In these tessellation artifacts, prints showed smooth surfaces with varying angles not aligned to the building direction which is attributed to the continuous printing in Xolography as opposed to many AM processes which suffer from layer-by-layer artifacts along building direction in these cases (Figure S2). Furthermore, while resolution of the negative feature in PEGDA/GelMA-hydrogels appeared to be limited to a range of 400 µm (Figure 3 F, Video S1), and UDMA prints showed a seemingly open 100 µm channel (Figure 3 H), PEGDA-hydrogels showed formation of a hollow channel with a diameter as low as 100 µm (Figure 3 G). The larger processing window for PEGDA-hydrogels allowed for fine tuning of the printing speed as a variable known to influence print resolution in photopolymerization AM. The smallest size of both positive and negative features herein was therefore found to be comparable to other rapid volumetric printing processes with hydrogel-based materials (∼20 µm and ∼100 µm, respectively) whereas the edges are more refined.^[11,12]^

### 2.3 Variation of stiffness can be achieved by tuning the Xolography process parameters

There are numerous mechanical properties of relevance for tissue engineering, however, we herein focused on the compressive modulus of the material to elucidate the dependency of stiffness of printed hydrogels on processing parameters (Figure 3 I). Besides being an important measure for selection of materials in tissue engineering, stiffness provides indications about the DoC as influenced by processing settings, specifically curing time and UV exposure.^[32, 33]^ We confirmed for hydrogel-based photoresins, that a higher conversion led to a higher macroscopic apparent stiffness on average (Figure S4, Figure S5): A GelMA/PEGDA hydrogel with a conversion of 44.5 % obtained a 87.2 % lower average stiffness than a post-processed comparison (59.4 kPa). A PEGDA hydrogel showed a decline in average stiffness of 54.4 % (2.3 MPa) compared to a post-processed state.

Stress-strain curves of printed and tested photoresins displayed a linear elastic region until a strain of 5 % (Figure 3 J–L). The observed stiffness variation on the order of three magnitudes between different photoresins was expected and likely corresponding to mesh-size and polymer concentration: GelMA/PEGDA-hydrogels displayed low stiffness in the range of 2.0–38.3 kPa (Figure 3 J).^[34, 35]^ Conversely, PEGDA hydrogels showed stiffness in the range of 0.5–6.5 MPa (Figure 3 K) and specimens from acrylate resins exhibited a stiffness in the range of 0.3–9 MPa (Figure 3 L). To gain perspective, the obtained range for GelMA/PEGDA-hydrogels was found to align with stiffness in soft living tissue including brain, kidney, and muscle tissue, while PEGDA-hydrogels in turn demonstrated stiffness close to the transition from soft to stiff living tissue, represented by gut, nerve, and cartilage tissue.^[36]^ This highlights the broad applicability of both investigated photoresins and Xolography as a suitable process for soft tissue engineering applications.

Beyond this broad applicability, we found that a prediction of the stiffness as a function of printing speed and UV intensity in Xolography is partially possible by using a central composite DoE (Figure 3 M) in combination with response surface methodology (Figure 3 N– P). For GelMA/PEGDA-hydrogels, 63.1 % of the variation in stiffness could be computed by an expression including the main effects of speed and intensity, as well as their first order interaction effect (Table S2, Table S3, Table S4). The direction of the main effects portrayed their impact on energy input and consequently photopolymerization: speed strongly influences stiffness negatively (p = 0.0009), intensity on the contrary is positively correlated with stiffness but its effect was not significant (p = 0.1). Even though the effect of intensity showed no significance, the interaction effect between speed and intensity was significant (p = 0.02). The minor influence of changes in speed at low intensities could be explained by a low share of activated isomers if irradiation was insufficient. At high intensities, an increased number of activated isomers was present and exposure time determined how many of these activated isomers participated in radical formation, resulting in increased crosslinking and stiffness (Figure 3 N). The high dependency of the GelMA/PEGDA-formulation on printing speed as the parameter, which is determining exposure time, might be attributed to the above-mentioned low amount of methacrylamide groups in combination with high MW. Considering the relatively large portion of stiffness variation that cannot be explained by the model (36.9 %), additional parameters such as temperature and viscosity of the photoresin might affect the process and should be considered in future regression models such as this one. Although GelMA/PEGDA hydrogels could be handled after printing, prints were soft and required post-processing with additional UV exposure. We performed the same study with a reduced irradiation time of 5 min to identify whether this post-processing step override effects from the printing process. The range of compressive modulus was reduced to 0.2–12.1 kPa. The model showed increased predictability (R^²^_adj_ = 81.1 %) and confirmed the hypothesis that prolonged post-curing diminish the effect of process parameters.

For PEGDA hydrogels, 67.2 % of the variation in stiffness could be explained by employing only the main effects of speed and intensity and deducing a linear relationship which pointed towards a direct linear dependency of compressive stiffness development on energy dosage (Figure 3 O). Both effects were highly significant (p = 0.0000, p = 0.0004, respectively) with a change in speed having a 6-times higher impact than a change in intensity on stiffness.

Similarly, UDMA printing showed a significant effect of both main parameters for pristine samples and could be predicted linearly (R^²^_adj_ = 76.2 %). As soon as post-curing was conducted, the processing parameters intensity and speed were not sufficient to predict the stiffness and all 2^nd^ order regression models solely based on these parameters and their interactions exhibited a lack of fit. Post-processing therefore has an effect on stiffness that masks the initially observed effect of the printing parameters, similarly to the trend observed for GelMA/PEGDA samples. The post-curing procedure of UDMA-based samples includes both additional UV-irradiation as well as a thermal curing cycle, as suggested by previous literature. The individual effects of the post-processing parameters UV intensity, exposure time, temperature, and time of the thermal cycle could be fruitful investigations for optimization of the UDMA-system.

Speed notably influenced mechanical properties of printed specimens for all the tested photoresins. This was likely due to its impact on the DCPI both on activation as well as on radical formation. UV intensity, in contrary, mainly affected its activation. Furthermore, PEGDA which showed a large processing window showed to be available for fine tuning of properties like feature resolution as described above. Considering the results of the mechanical study and high resolution of the process, we hypothesized that the processing parameters can be utilized to locally control material stiffness within a printed part.

## 3. Significant modulation of stiffness can be implemented by grayscaled projection on microscale

The impact of spatially tunable mechanical cues on cell behavior, including cell alignment and anisotropic migration, has been demonstrated by modulated stiffness regions in volumetric technologies like FLight, although primarily on a single plane.^[14]^ Yet, higher control over local mechanical properties in 3D is still required to achieve a fully engineered microenvironment within printed constructs. Grayscale or gray-tone projection has recently emerged as a promising approach to unlock such mechanical control in 3D both in elastomers and hydrogels; however, so far it has been limited to non-volumetric techniques such as digital light processing (DLP) that generally involve longer processing times, lower throughput, and thus lower compatibility with cell-laden printing.^[37, 38]^ Interestingly, the grayscale strategy can be exploited in Xolography, since the linear relationship between stiffness and processing parameters for PEGDA photoresins (Figure 3) can be utilized to modulate stiffness throughout a print by process adjustments with a single photoresin formulation *in situ*. Freedom of the stiffness manipulation was, however, restricted to the printing direction if energy input was modulated based on printing speed and UV intensity, as is common in FLight. To navigate stiffness modulation within a 3D part, we continued with a grayscaled projection image of the visible light source. The visible light projection in combination with the movement of the cuvette in z-direction theoretically allowed for full control over stiffness down to a voxel defined by the projected pixel size of 4.6 µm x 4.6 µm (Figure 4 A). This pixel size is defined by the projector’s 4K resolution (3840 x 2160 pixels) and the focused projection width of 10 mm and height of 17.7 mm inside the cuvette. By employing a grayscaled projection, the energy input for the radical photopolymerization reaction was manipulated (Table S5), as opposed to stiffness modulation via process parameters which targeted the activation reaction of the DCPI. To investigate the capability and limitations of Xolography to adjust the stiffness of a part within a single-cure print, we studied three cases in which varying stiffness is realized on a global (1) and local scale (2), as well as within a continuous (3) gradient (Figure 4 A). For improved readability, results are presented referring to the implemented RGB values within the manuscript. Corresponding intensity of the projected light is detailed in the legend of Figure 4 B and supplementary information (Table S4).

**Figure 4.**
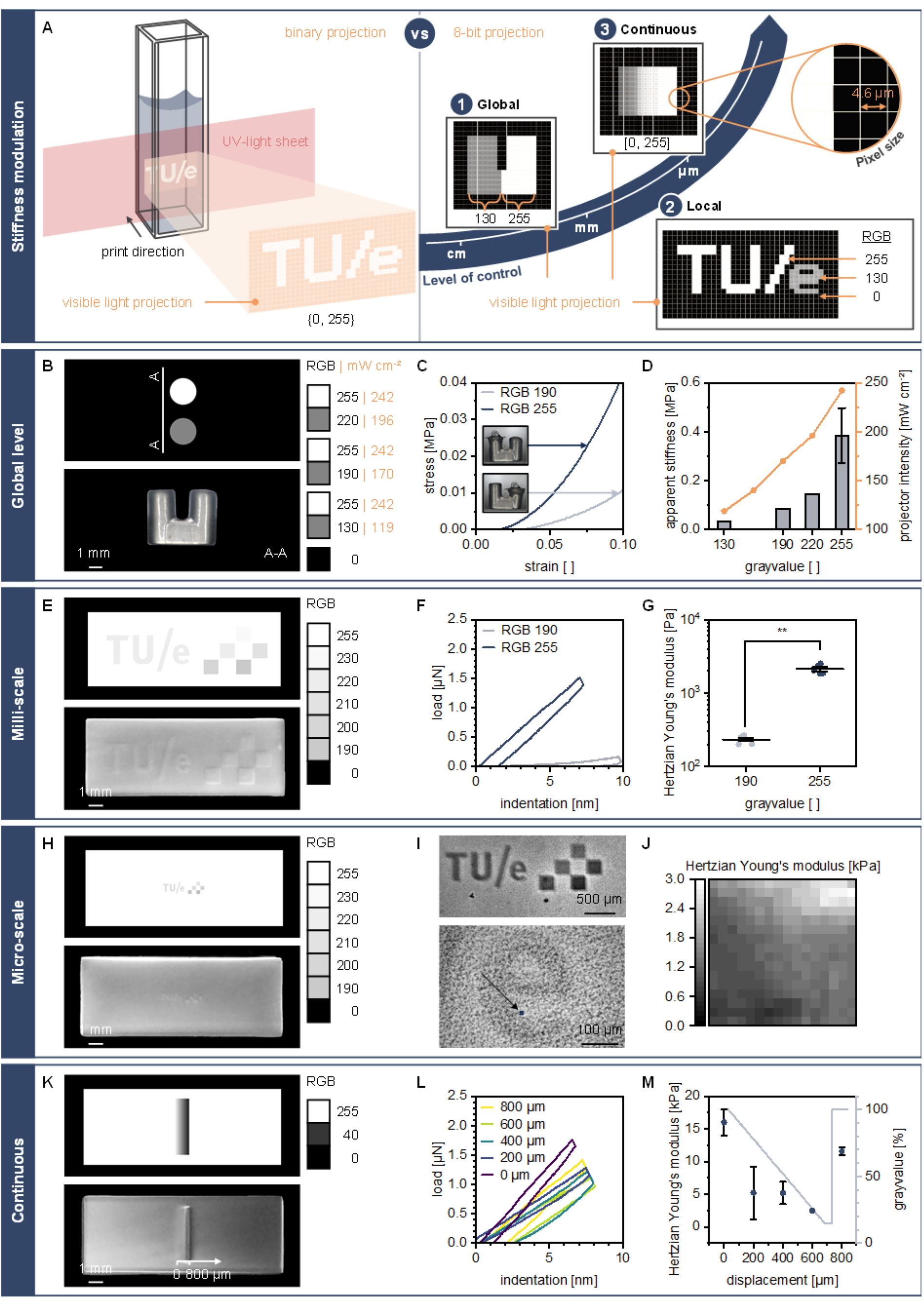
Xolography demonstrates the potential for local stiffness modulation within a single print and in a single step. A) Schematic principle of the traditional binary projection and the adjusted grayscaled projection and a simplified representation of the employed specimens. B) Grayscaled projection mask and printed cylinders for compression testing. C) Representative stress-strain curve for a part printed with RGB = 255 (left side) and RGB = 130 (right side), n=1. D) Compressive stiffness of parts printed at different RGB-values and the corresponding actual intensity of the projector, n = 1 (n = 6 for RGB = 255, depicted are mean ± SD). E) Grayscaled logo with sections of different RGB-values. F) Nanoindentation of two regions. G) Hertzian Young’s modulus of the tested regions, n = 5, two-sided Mann-Whitney U test, p ≤ 0.008, mean ± SE shown. H) Grayscaled logo with sections of different RGB-values on microscale. I) Closeup of the printed text via brightfield (top: 20x objective, bottom 2.5x objective). J) Hertzian Young’s modulus mapping of the region indicated in I, n = 1. K) Continuous gradient with 600 µm in width. L) Nanoindentation at distinct locations of the gradient, n = 1. M) Derived stiffness values and corresponding grayscaled percentage for the gradient, with an estimated gradient of 20 Pa µm^-1^, n = 3, visualization of mean ± SD. Printing speed = 5 mm min^-1^ for all samples.

Global stiffness differences were successfully implemented within a print of two connected cylinders (Figure 4 B). The cylinder printed at full exposure (RGB 255) showed a higher compressive stiffness around 400 ± 100 kPa compared to the cylinder printed at a grayscaled exposure (RGB 130, 190, or 220) with 33, 87, and 147 kPa, respectively. Moreover, measurements of the actual intensity of the projector showed the same trend as the stiffness values for increasing RGB-values (Figure 4 C, D).

We furthermore showed the transfer to local stiffness modulation on milli-scale level with plates containing multiple RGB-value regions (Figure 4 E). Notably, material heterogeneity was visible on the printed part itself, showing differences in refraction between the base plate (RGB 255) and small areas with adjusted grayscale. We selected the region of RGB 190 for indentation measurements as an example. We observed a highly significant difference (p ≤ 0.008) in the indentation curves (Figure 4 F) and the derived stiffness values (Figure 4 G): an average stiffness of 2100 ± 300 Pa was measured for regions with full exposure and 230 ± 30 Pa for regions with reduced exposure (RGB 190). We demonstrated that a decrease of 25 % in grayvalue resulted in a lower stiffness. To determine whether grayscale-printed hydrogels undergo changes in stiffness contrast across regions with high and low gray value over time, we tested RGB 190 and 255 regions in samples after prolonged cold storage. We observed that grayscale-printed hydrogels do not exhibit significant changes in Hertzian Young’s modulus between 2 and 8 weeks of storage for RGB 190 and RGB 255 (Figure S6). However, the contrast in stiffness, defined as the ratio between the Hertzian moduli of low and high projector intensity regions, was significantly reduced (p ≤ 0.035). The Hertzian modulus for RGB 255 prints was approximately an order of magnitude larger than for RGB 190 prints even after 8 weeks, indicating that, despite a contrast decay, the final contrast value remains considerable after prolonged storage.

We further leveraged the projector’s resolution to decrease the scale of stiffness modulation to the microscale (Figure 4 H). The interface between full exposure (RGB 255) and reduced exposure (RGB 220) regions was mapped to assess the presence of stiffness differences (Figure 4 I). Over a total scan area of 5 µm x 5 µm, a trend in stiffness was found to follow the grayscaled projection. The scanned area closer to the RGB 255 region showed up to five times higher stiffness than that of the RGB 220 region (Figure 4 J, Figure S7). While the multiple testing techniques used are not directly compatible for comparison of the absolute stiffness values among global, milli-, and microscale, key differences in the stiffness magnitude were found and thus inform the advancement of constructs with spatially controlled mechanical properties.

We finally implemented a continuous gradient (Figure 4 K) and were able to show a continuous decrease in stiffness with decreasing grayvalue (Figure 4 L, M). We therefore demonstrated the possibility to employ visible light projection with a simple grayscale to modulate the stiffness within a single print, observing a measured stiffness gradient of about 20 Pa µm^-1^. While stiffness gradients have been shown before for tomographic volumetric printing, this is the first time that one-step stiffness modulation in volumetric printing of a single hydrogel is shown, to the best of our knowledge.^[13]^ Considering the potential for cell guidance based on the geometrically restricted FLight process, this approach unlocks new opportunities for full 3D control over stiffness with applications in tissue engineering and advanced *in vitro* modeling.^[39]^

## 4. 4D volumetric printing of thermoresponsive hydrogels allows for controlled shape change

Given the potential for locally modulated DoC using grayscaled projections, we implemented the 4D fabrication of stimuli-responsive hydrogels to generate constructs with reversible spatiotemporal control over shape (Figure 5 A). We selected NIPAAm, a commonly used thermoresponsive acrylamide-based monomer that transitions from hydrophilic to hydrophobic state at the lower critical transition temperature (LCST) of 32 °C, which enables shrinking behavior of NIPAAm hydrogels above this temperature by expulsion of trapped water molecules. NIPAAm was combined with GelMA as a macromer, which provides a) gelation of the hydrogel precursor at room temperature and b) a source of branching points to allow for crosslinking without the need for other branching monomers that tune the hydrogel network density, such as the commonly used *N*,*N*-methylenebisacrylamide (MBA).^[40]^ Crosslinking was feasible with a formulation of NIPAAm and GelMA at a 2:1 weight ratio, although heterogeneous curing was observed, showing un-crosslinked photoresin at the center of the printing region (Figure 5 B). When using similar BIS-TRIS concentrations than for GelMA/PEGDA-based photoresins, higher energy dosage was then required, which made overcuring more likely. Therefore, the BIS-TRIS concentration was increased to 2 M to enhance printability in the same parameter range (Figure 5 B). The increased availability of BIS-TRIS facilitated initiation of the radical crosslinking reaction, which allowed for the fabrication of centimeter-scale thermoshrinking NIPAAm/GelMA hydrogels with complex structures, such as gyroids (Figure 5 C, Figure S8, Video S2).

**Figure 5.**
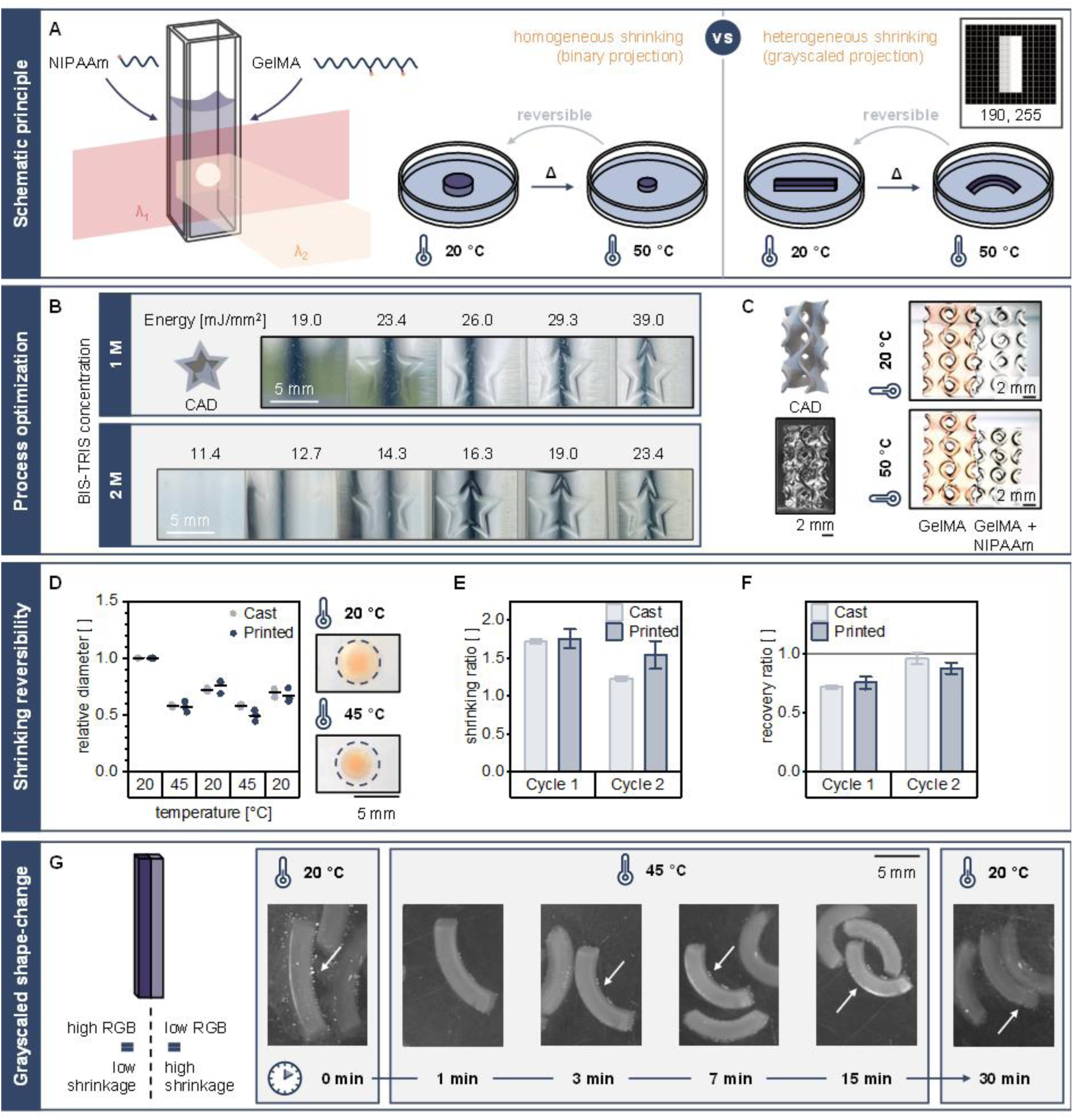
4D fabrication of thermoresponsive complex 3D structures with shape change beyond isometric shrinkage. (A) Schematic principle of binary and grayscaled Xolography for 4D printing of NIPAAm-based hydrogels. (B) Effect of BIS-TRIS concentration on the processing window (energy dosage) of NIPAAm/GelMA hydrogels. (C) Complex gyroid structures based on optimized photoresin formulations and their thermoshrinking behavior. (D) Effect of thermal cycling on NIPAAm/GelMA hydrogel size (n = 3, mean). (E–F) Shrinking capacity and plasticity after thermal cycling of NIPAAm/GelMA hydrogels (n = 3, mean ± SD). (G) Thermoresponsive bending of Janus hydrogel beams obtained by grayscaled Xolography.

Due to time-dependence of the thermoshrinking response, we proceeded to evaluate the shrinking kinetics and reversibility of printed NIPAAm/GelMA hydrogels with a sol fraction of 62.2 ± 0.4 %. Shrinking at 45 °C was found to be reversible at comparable rates for both printed and cast hydrogels, though both groups showed plasticity upon re-swelling at 20 °C (Figure 5 D). Moreover, thermal cycling deteriorated the shrinking ability to a larger degree for cast hydrogels than for printed hydrogels, since the average shrinking ratio of cast and printed hydrogels decreased by 28% and 12% after the first cycle, respectively (Figure 5 E). Additionally, both cast and printed hydrogels showed a lower ability to recover after the first shrinking cycle than after the second, as observed through the recovery ratio (Figure 5 F). Overall, the shrinking plasticity of this hydrogel system can be ascribed to the presence of GelMA as macromer crosslinker, which might amplify the viscous component of the hydrogel network and thus limits the recovery of the fully swollen state. Such irreversible behavior under thermal cycling has been previously shown to be less prominent in NIPAAm photoresins containing MBA as crosslinker.^[37, 40]^ However, these hydrogel systems are more commonly used in lithography-based 3D printing approaches that do not require considerable viscosity of the photoresin to ensure print quality, and thus at present, volumetric printing techniques such as Xolography still rely on rheological modifiers such as GelMA.

Based on the ability of grayscaled projections to locally control the DoC, we developed a strategy to spatiotemporally program hydrogel shape beyond isometric shrinking. Considering that swelling and shrinking are dependent on the DoC of NIPAAm-based hydrogels, we used grayscaled Xolography to produce Janus hydrogels –integrating two distinct material properties separated by an interface– in a one-step process that would enable shape change such as bending or buckling.^[41]^ To test this strategy, Janus beams were fabricated by using two grayvalues to create a high DoC and a low DoC region separated by an axially oriented interface (Figure 5 G). Upon heating, the hydrogel beams underwent bending due to differential shrinkage, which remained partially reversible upon cooling, similarly to hydrogels printed using binary projection (Video S3). Evidence from this proof-of-concept suggests that grayscaled Xolography in combination with stimuli-responsive hydrogels has the potential to generate constructs with tunable spatiotemporal control over shape change, especially by leveraging the micrometer scale of pixel size in Xolography. Previously, the emergence of bending in Janus or multi-region beams or films has been exploited via multi-cure or sequential fabrication processes to introduce shape change behavior into 3D-printed biomaterials.^[42, 43]^ Only recently have single-cure grayscaled projections been applied in the context of 4D printing, primarily in TPP and DLP techniques, in which throughput is limited.^[37, 38, 44]^ Thus, grayscaled Xolography of stimuli-responsive hydrogels is a single-step strategy to create spatiotemporally programmable centimeter-scale biomaterial platforms with promising implications for the fields of soft (bio)robotics, tissue engineering, and morphogenesis modelling.

## 5. Aggregate bioprinting shows cytocompatibility and confirms the need for local control of mechanical properties

Cell-laden printing, or bioprinting, is of crucial value to move from uniform 3D cell cultures towards tissue-mimicking spatial organization and differentiation of cells. Xolography has great advantages for cell-laden printing because it has short processing times, maintains sterility in an enclosed environment and does not subject cells to shear stresses during printing. We therefore developed a pipeline for cell-laden printing, using the newly developed hydrogel formulations. We first assessed cytocompatibility of the individual components of the photoresins with multipotent human mesenchymal stromal cells (hMSCs) in cast hydrogel discs (Figure 6 A). Metabolic activity of hMSCs in hydrogels that contained either DCPI or BIS-TRIS was reduced compared to a 10 % w/v GelMA hydrogel control at one and seven days after embedding, indicating a reduced cell viability and limited cytocompatibility of the DCPI and high concentrations of the coinitiator (BIS-TRIS) in their unreacted form. This highlights the importance of minimizing exposure time of the cells to the unreacted forms of both DCPI and coinitiator prior to crosslinking, which occurs during printing. After printing, further exposure should be minimized through thorough washing procedures to remove unreacted DCPI and coinitiator after crosslinking. Large variation between technical repetitions in DCPI samples were likely due to processing time, further supporting the importance of limiting exposure to the DCPI system.

**Figure 6.**
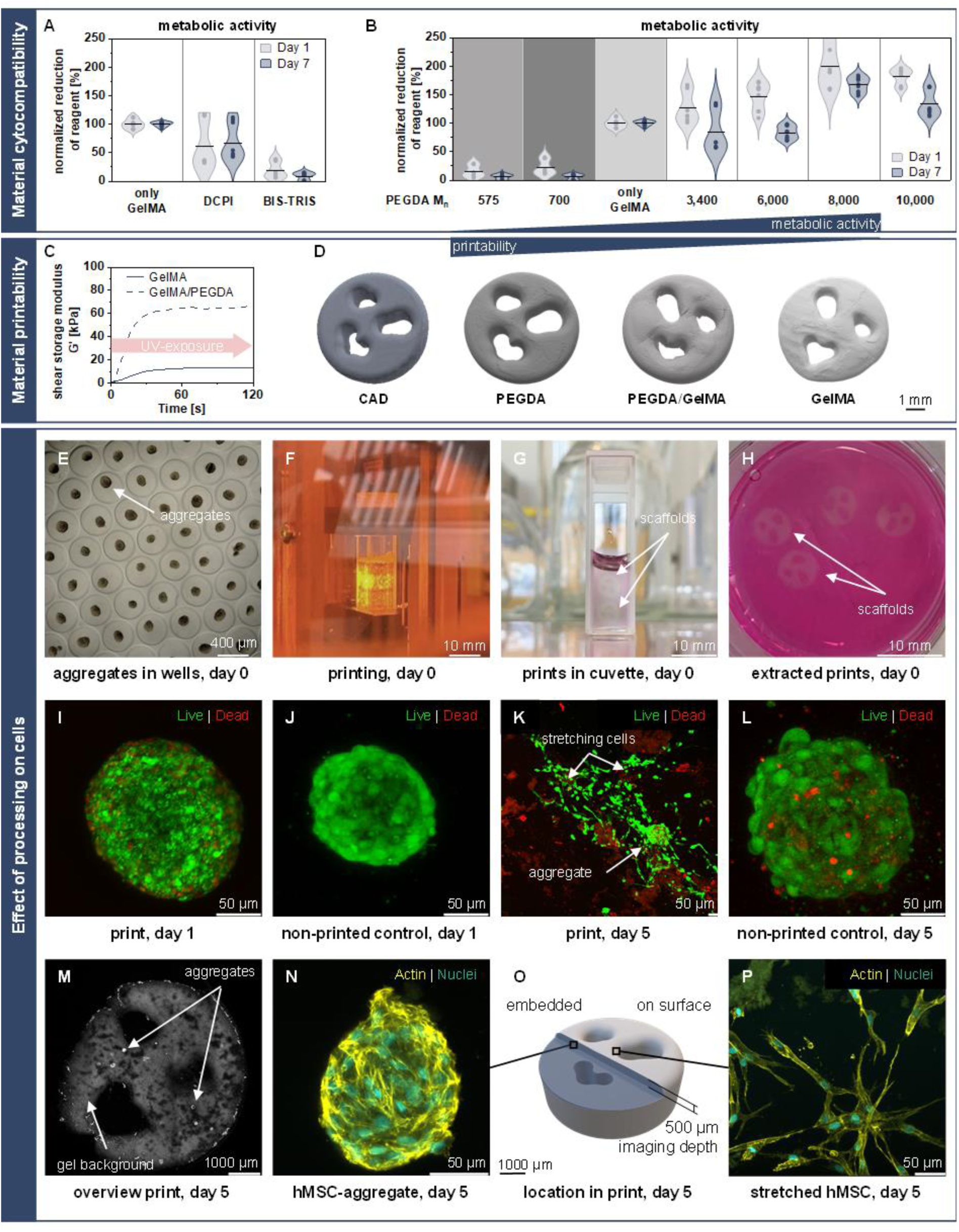
Adaptation of the material formulation shows biocompatibility and potential to print cell-laden constructs. (A) Influence of DCPI and BIS-TRIS on metabolic activity (Alamar Blue) of hMSCs in cast GelMA (10%), n = 9 samples from 2 independent experiments, mean reported. (B) Influence of M_n_ in PEGDA on metabolic activity (Alamar Blue) of hMSCs in cast GelMA/PEGDA (10% / 1.5 % respectively), n = 9 samples from 2 independent experiments, mean reported. (C) Crosslinking behavior of hybrid and synthetic photoresins upon UV irradiation. (D) MicroCT scans of printed scaffolds based on photoresin composition, n = 1. (E) Formation of hMSC-aggregates in a microwell-plate. (F) Printing of an aggregate-laden hydrogel. (G) Printed constructs in a cuvette after printing. (H) Scaffolds after post-processing in culture medium. (I-L) Live/dead staining (Calcein-AM (green)/Ethidium homodimer (red)) of (I) a printed hMSC-aggregate in a GelMA hydrogel on day 1 after printing, (J) non-printed control aggregate in its microwell on day 1, (K) a printed hMSC-aggregate in a GelMA photoresin on day 5 after printing and (L) non-printed control aggregate in its microwell on day 5. (M) Overview of printed construct with hMSC-aggregates. (N) Morphology of an incorporated hMSC-aggregate showing phalloidin-stained actin (yellow) and DAPI-stained nuclei (aqua). (O) Schematic representation of different localization of morphologically different cell-aggregates (N, P) in a print. (P) hMSC-aggregate close to the surface.

Printing hMSCs in the hybrid photoresin with 10 % w/v GelMA and 10 % w/v PEGDA that was successfully used for printing in Section 2.2, did not yield viable samples and reducing PEGDA to 2.5 % w/v and 1.5 % w/v did not clearly improve metabolic activity (data not shown). Based on previous research of hydrogels with high PEGDA concentrations, we hypothesized that the low M_n_ = 575 of the PEGDA influenced cell viability and increasing the M_n_ of PEGDA would result in increased cell metabolic activity.^[45]^ Indeed, metabolic activity of hMSCs in cast disks increased when substituting PEGDA with low M_n_ with PEGDA with higher M_n_, indicating increasing viability with increasing M_n_ up to a M_n_ = 8,000 (Figure 6 B). Hybrid gels containing PEGDA with M_n_ > 3,400 even outperformed hydrogels containing only GelMA-hydrogels.

Addition of low M_n_ PEGDA to the hydrogel formulation improved hydrogel printability as reported previously, which is likely due to faster crosslinking kinetics (Figure 6 C).^[46]^ This results in a wider printing window (Figure 3 B) and a closer representation of the CAD geometry (Figure 6 D). However, the addition of high M_n_ PEGDA led to no noticeable improvements in printability or geometrical representation, but instead resulted in an increase of overcuring and thus poorly defined prints. Although high M_n_ PEDGA improved cell viability, the resulting loss in printability showed a trade-off between print resolution and cell viability for the hybrid polymer formulation. We therefore decided to use a pure GelMA-hydrogel to assess the suitability of the printing process for bioprinting. Advancement of the resolution for these gels could be reached by adjusting the photoresin refractive index to match with the refractive index of cell aggregates as described in literature.^[11]^

Pre-formed hMSC aggregates (initially 500 cells, Figure 6 E) were printed in sterile GelMA-hydrogel to assess cytocompatibility of the printing process. Printing with hMSCs in aggregates kept individual cells within close distance to each other to promote essential cell-cell interactions and delivered them in a protected manner. The aggregates remained evenly distributed due to hydrogel gelation at room temperature, and aggregate-laden scaffold geometries were printed, washed and cultured over five days (Figure 6 F, G, and H). Printed constructs resembled the CAD design (background signal staining) and incorporated hMSC-aggregates were randomly distributed (Figure 6 M). Importantly, printed cell aggregates showed viable cells present after one and five days in culture, visualized with Live/Dead staining (Figure 6 I, K) and were compared to non-printed controls (Figure 6 J, L). One day after printing, dead cells occurred mainly on the outside of the aggregate, likely due to previously described cytotoxic effects of the material or disturbance by handling and UV exposure (Figure 6 I). However, a large fraction of the cells in each aggregate maintained their viability also five days after printing (Figure 6 K), and cells started to migrate out of the aggregates and spread over the hydrogel surface.

Interestingly, cells behaved differently depending on the location in the hydrogel scaffold (Figure 6 O). The migrating cells transitioned to the typical morphology of stretched hMSCs on culture plate surfaces and were mainly observed for aggregates close to the surface of the print (Figure 6 P), while aggregates which were located at a distance from the surface largely remained in their initial spherical shape (Figure 6 N). Whether this difference in behavior is due to geometry, material stiffness or nutrient/oxygen gradients remained unclear. Nevertheless, this highlights the importance of spatial control over geometry and material stiffness to actively influence the cell or aggregate microenvironment to guide cell fate.^[3]^ Additionally, it raised the urge for the controlled placement of cells in materials during photopolymerization-based printing processes.

This study also shows that the rapid crosslinking during Xolography minimizes cell exposure to unreacted groups, preventing harmful effects. While 100 % conversion is not achieved after printing, as confirmed by FTIR data (Figure S5 A–B), unreacted groups do not noticeably affect cell viability, likely due to effective removal during washing and post-curing. Overall, we provided the first proof that the Xolography printing process can be used for cell-laden printing, and future studies should further optimize hydrogel formulations to further enhance cytocompatibility.

## 6. Conclusion

We established Xolography as a technology to create centimeter-scale hydrogel geometries with structural resolution on cell-level and locally controlled stiffness on microscale within minutes. Achievable feature resolution is different for naturally-derived, synthetic, and hybrid hydrogels, and is dependent on MW, functional groups, polymer content, and thermal decay of the specific photoinitiator system. The resolution achieved herein is in the order of magnitude of commonly used tomographic volumetric printing processes but realizes more pronounced edges. Hybrid and synthetic hydrogels achieved higher resolution of negative features than naturally derived hydrogels, but only the latter were compatible with cell-laden printing. Measures to further improve the resolution in bioprinting include tuning of optical properties, refinement of biocompatible hydrogel formulations, and the continued development of the DCPI.^[11]^ While we manufactured centimeter-scale hydrogel constructs (height = 16 mm, width = 8 mm) herein, an increase of the available build volume for printed hydrogels will further support an extension of the range of applications.

The compressive stiffness of hydrogel formulations was tunable within a specific processing window through the parameters printing speed and UV intensity, which determine the applied energy. Inspired by this knowledge, we investigated whether stiffness can also be modulated by changing visible light intensity, thus modulating the secondary step of the photopolymerization reaction. While grayscaling is used in 2D photopatterning strategies, here grayscaling of the visible light projection enabled local stiffness modulation along all three axes. Xolography enables this modulation by independent control of energy input per voxel and restriction of visible light-induced polymerization to a defined UV sheet. Other volumetric printing processes like tomographic printing in turn link polymerization degree across all voxels in a plane via mathematical transformation which makes independent modulation physically or algorithmically unfeasible for certain patterns. Stiffness modulation of thermo-responsive hydrogels further permitted us to incorporate shape changes in prints and thereby to add a reversible dynamic response, with a wide range of potential applications in tissue engineering, advanced in vitro models, and soft (bio)robotics.^[5]^

A promising next step is the use of Xolography to print (living) constructs with features that allow us to study the impact of local stiffness, scaffold degradability, and eventually viscoelasticity on cell morphology and behavior. These studies are essential to enhance representation of the natural extracellular environments for in-vitro models and to develop instructive scaffolds that can modulate cell behavior or even control cell fate. The combination of grayscaled and cell-laden printing with in-line monitoring of cell location and Xolography in flow might further unlock precision and scaling in tissue engineering.^[16, 18]^

## 7. Experimental section

### Characterization of the DCPI and coinitiator

Absorbance spectra of the DCPI and its thermal relaxation behavior in the presence of different coinitiators were measured as described previously.^[17]^ Absorbance measurements of the spiropyran were conducted without additional irradiation, while the measurements of merocyanine were conducted under orthogonal UV excitation (375 nm). Half-life was determined by initially exciting the DCPI and measuring the decay in absorbance upon interruption of irradiation. A biexponential model was fitted to the curve in order to determine the values for the shorter and longer half-life values.

### Photoresin preparation and printing of cell-free constructs

Methacrylated gelatin (GelMA) was prepared following a previously described protocol.^[47]^ Unless stated otherwise, all materials were obtained from Sigma Aldrich (USA). 10 g of Type A gelatin from porcine skin with a bloom of 300 g was dissolved in 100 ml of ultra-pure water (UPW) and functionalized with 6 g of methacrylic anhydride in the dark at 50 °C for three hours, followed by dialysis against UPW in the dark at 40 °C for four days. Water was changed twice per day.

The DoM was determined with proton nuclear magnetic resonance (^1^H NMR).^[48]^ 5 mg of GelMA or gelatin were dissolved in 800 µl deuterium oxide (D_2_O). Measurements were conducted on a Bruker Avance 400 MHz NMR spectrometer (Bruker Coorporation, USA. Data was processed with MestReNova (Mestrelab Research S.L., Spain). Phase correction was conducted manually, and spectra were baseline corrected (Figure S9). Data were normalized to the phenylalanine signals at 7.2–7.5 ppm. Reduction of the lysine amino acids at 2.8–2.95 ppm was determined by integration and the DoM is determined by the following equation:

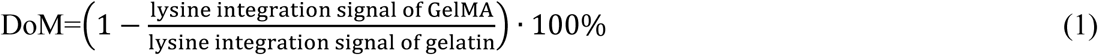

Hybrid and naturally-derived hydrogel solutions were prepared by dissolving GelMA at 10 % w/v in UPW at 37 °C overnight, shielded from light. Unless indicated otherwise, BIS-TRIS was added to a final concentration of 800 mM. For semi-synthetic hydrogels 10 % w/v PEGDA (M_n_ = 575) was added. Temperature was raised to 45 °C and mixing was enhanced by alternating between a rolling plate and water bath. DCPI XC-577 (available as DCPI 5001, xolo GmbH, Germany) was added for to a final concentration of 0.015 % w/v. 1.5 mL of photoresin was filled in single-use, UV-transparent fluorescence cuvettes (Carl Roth GmbH, Germany) and centrifuged at 300 x *G* for 2 min to remove potential air bubbles. Cuvettes were allowed to gelate in 4 °C storage for 30 min.

Synthetic hydrogel solutions were prepared by mixing DCPI-containing PEGDA with a commercially available aqueous solution of TEOA and a polyacrylic acid for rheology control. Under constant stirring, 24.0 mg DCPI XC-471 (available as DCPI 3001^[49]^, xolo GmbH, Germany) dissolved in 40 g PEGDA (M_n_ = 700) was added dropwise via a syringe to 60 g of xoloPEGDA_Component-A (xolo GmbH, Germany) to result in a homogeneous formulation. UDMA-based acrylate resin was used as benchmark material and was obtained as a pre-mixed solution under the commercial name xoloOne (xolo GmbH, Germany).

For thermoshrinking hydrogels, precursor solutions were prepared containing final concentrations of 10 % w/v NIPAAm, 5 % w/v GelMA, 0.015 % DCPI (dissolved in PEGDA M_n_ = 575 at 1 wt %), and 2 M BIS-TRIS at pH = 7.4 as coinitiator in UPW. Control cast thermoshrinking hydrogel disks (height = 2 mm, diameter = 5 mm) were prepared containing final concentrations of 10 % w/v NIPAAm, 5 % w/v GelMA, and 0.1% w/v lithium phenyl-2,4,6-trimethylbenzoylphosphinate (LAP) as photoinitiator in UPW, followed by UV exposure for 5 min.

Xolography was performed using a customized light-sheet volumetric printer based on the setup Xube (xolo GmbH, Germany), equipped with two UV laser diodes (375 nm) and a projector (3840 x 2160 pixels). Prints of synthetic hydrogels were conducted on a commercially available 405 nm version Xube printer (xolo GmbH, Germany).

Uncured photoresin was re-dissolved at 37 °C and parts were washed in prewarmed phosphate-buffered saline (PBS) at 37 °C. GelMA, GelMA/PEGDA, and NIPAAm/GelMA hydrogels that were obtained from binary projections were post-cured in PBS with 0.1 w/v % LAP for 5 or 10 min within the Asiga Flash UV curing lamp (Asiga, Australia). An increase in curing time or LAP concentration did not decrease the sol fraction of the prints (Figure S10). UDMA-based prints were post-processed as previously described.^[17]^

### Characterization of printed hydrogel-based constructs

Mechanical characterization was performed via compression tests conducted in accordance with ASTM D695–15.^[50]^ Cylindrical parts were printed with a height of 4 mm and a diameter of 2 mm. Unconfined compression was conducted in air at a speed of 1.3 mm min^-1^. Compression testing of hydrogels was performed with a MicroTester G2 (CellScale Biomaterials Testing, Canada), utilizing a circular tungsten beam (modulus = 411 GPa) with diameter of 1.016 mm for synthetic hydrogels and 0.4064 mm for GelMA-containing hydrogels. Acrylate resins were compression tested with a MTS Criterion Model 42 (MTS Systems Corporation, USA) and a 50 N s-beam load cell. Load/displacement curves were obtained from both devices with an initial pre-displacement of 1 µm. Data was transformed in engineering stress/strain curves using the dimensions of the specimen prior to testing which were obtained by measurements with Fiji.^[51]^ Compressive stiffness was deducted by fitting a linear regression to the range of 1–5 % of strain. Data analysis was performed with Python in PyCharm 2022.2.1 (JetBrains s.r.o., Czech Republic). Sol fraction was determined from printed post-cured hydrogel disks, which were weighed for their initial wet weight (m_wet,t=0_) and lyophilized to obtain their initial dry weights (m_dry,t=0_), or incubated in PBS at 37 °C for 24 h, weighed (m_wet,t=1_), freeze-dried and re-weighed (m_dry,t=1_). The sol fraction was determined as follows:

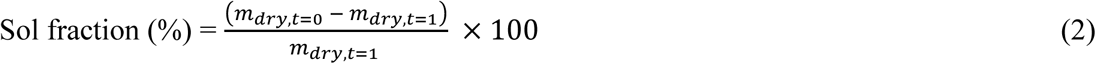

Fourier-transform infrared spectroscopy (FTIR) was performed on uncured solutions, printed hydrogels, and printed and post-cured hydrogels using an IRTracer-100 spectrophotometer (Shimadzu, Japan) coupled with an attenuated total reflection (ATR) accessory, collecting an average of 64 scans with 1 cm^-1^ resolution in the 800–4000 cm^-1^ range. The acrylate C–H scissoring band (1390–1440 cm ^-1^) was used to monitor unreacted double bonds, and the peak area was normalized using the carbonyl band (1710–1758 cm ^-1^) for PEGDA and the amide II band (1600–1700 cm ^-1^) for GelMA/PEGDA as internal reference. Hydrogel precursor conversion was estimated from the normalized acrylate peak area (A) of printed, uncured, and post-cured samples using the following equation:

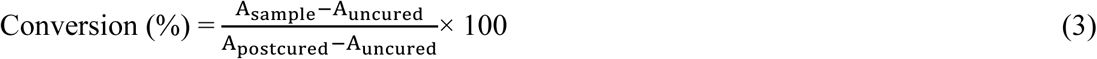

Local stiffness of grayscale-printed PEGDA hydrogels after 2 weeks and 8 weeks of storage at 4 °C was assessed at room temperature in PBS using a Piuma nanoindenter (Optics 11, the Netherlands), using a probe with a spherical tip radius of 28 µm and a spring constant of 0.51 N m^-1^. For each grayvalue region, at least 3 randomly selected sites in each of 5 different samples were tested. Effective elastic modulus (stiffness) was estimated from force-indentation curves by fitting a Hertzian contact model given by the following equation:

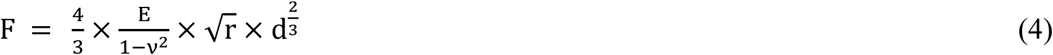

Where *F* is the applied force, *E* is the Young’s modulus, *v* is the Poisson’s ratio, *r* is the spherical tip radius, and *d* is the indentation depth.

Mechanical mapping of grayscale PEGDA prints was performed via atomic force microscopy (AFM) on a Cypher Environmental Scanner (Oxford Instruments, UK) controlled by Igor Pro software using a colloidal polymethyl methacrylate-based CP-CONT-PM probe (Nanosensors) with a spring constant of 0.13 N m^-1^ and a tip radius of 750 nm. The cantilever was calibrated using GetReal function in Igor Pro software. Sample and cantilever were submerged in PBS in a closed gas cell. Force curves were collected over a force distance of 4 µM at a scan rate of 0.1 Hz, using an absolute trigger point of 2 x 10^-9^ N. Force maps were acquired by collecting 16 x 16 curves in an area of 5 µm x 5 µm. Force curves were fitted to a Hertz model (Equation 4) using a tip Poisson’s ratio of 0.4 to extract the Young’s modulus.

The 3D shape of simple hydrogel prints without delicate features was assessed with a MicroCT 80 (Scano Medical AG, Switzerland) using a scan intensity of 88 A, energy of 45 kVp, integration time of 1000 ms and a voxel size of 11.4 µm. Samples were imaged in air in a sealed petri dish. Prior to imaging, hydrogel samples were soaked in a contrast agent containing 15 mg ml^-1^ iodine in PBS overnight. Images of printed samples were taken with a digital microscope (AD407 by Andonstar Technology Co., Ltd., China) with hydrogel samples immersed in water.

### Grayscaled print files

Grayscaled projections were created from a series of respective 3840 x 2160 pixels PNG files using the VideoCreator script (xolo GmbH, Germany) and imported in the Xube software. Using the standard settings for customized print file import, each PNG image file resembled a 5 µm thick slice of the target object. Accordingly, repetitive batches of grayscaled images were created using RGB-defined filling of 2D shapes in Microsoft Paint. Image order for print file generation was indicated by an ascending number in respective file names.

### Characterization of thermoshrinking response

Grayscale-projected NIPAAm/GelMA hydrogels were tested with no post-curing treatment. Printed or control cast NIPAAm/GelMA hydrogels were kept in a water bath at 20 °C, followed by immersion in a water bath at 45 or 50 °C. In the case of cyclic testing, two heating and cooling cycles were performed. Samples were allowed to stabilize for 15 min for each run or cycle, and size and shape change were tracked throughout using a M5 camera (Basler AG, Germany) controlled with a MATLAB interface (MathWorks). Shrinking ratio was defined as the swollen-to-shrunken diameter ratio, and recovery ratio was defined as the reswollen-to-shrunken diameter ratio. Comparison to a non-thermoresponsive hydrogel was performed on gyroid structures (height = 15 mm, width = 7.5 mm). Hydrogel bending due to thermoshrinking was assessed on hydrogels with beam geometry (square cross-section, length = 15 mm, width = 3 mm) fabricated using grayscaled projections (RGB values of 190 and 255) to generate two axially oriented regions with different DoC. Beam curvature under heating over time was quantified using the Kappa Curvature analysis plugin in Fiji software.^[51]^ All shrinking tests were performed with a sample size of n = 3–4.

### Culture of cells and cell aggregates

hMSCs were isolated from human bone marrow of a healthy adult male donor (1M0125, Lonza, USA; collected under their institutional guidelines and with informed consent; experiments were performed at Eindhoven University of Technology according to Dutch occupational health legislation) and characterized as previously described.^[52]^ hMSCs were cryopreserved in fetal bovine serum (FBS, Serana Europe GmbH, Germany) containing 10 % dimethyl sulfoxide (DMSO) until use in experiments. Thawed hMSCs (passage 4–6) were collected in high glucose Dulbecco’s Modified Eagle’s Medium (DMEM) with L-glutamine (hg-DMEM, 41966, Gibco™) before seeding (2.5 x 10^3^ cells cm^-2^) and expansion until confluent (7–10 days) in hg-DMEM supplemented with 10 % FBS, 100 U ml^-1^ of Penicillin and 100 U ml^-1^ of streptomycin (P/S, Gibco™), 1x MEM Non-Essential Amino Acids (NEAA, Gibco™) and 1 ng ml^-1^ recombinant human basic fibroblast growth factor (bFGF, PeproTech®). When 80–100 % confluent, hMSCs were dissociated using 0.05 % Trypsin-EDTA (Gibco™) before inactivating the trypsin with medium containing 10 % FBS. Cells were pelleted (270 x *G*, 7 min), counted and resuspended either in PBS for casting or in aggregate medium for aggregate assembly (1 part DMEM/Nutrient Mixture F-12 Ham (F12, Gibco™) and 2 parts human endothelial SFM (Gibco™) supplemented with 100 U ml^-1^ P/S, 1 x NEAA, 0.1 mM 2-mercaptoethanol (50 mM, Gibco™), 1× N2 (Gibco™), 1× B27 supplement (Gibco™), 1 ng ml^-1^ bFGF, 20 ng ml^-1^ recombinant human epidermal growth factor (hEGF, Chimerigen Laboratories, Switzerland), 20 ng ml^-1^ recombinant human platelet derived growth factor (hPDGF-AB, PeproTech®), 20 ng ml^-1^ recombinant human oncostatin M (hOSM, Gibco™), 40 ng ml^-1^ recombinant human insulin-like growth factor 1 (hIGF-1, PeproTech®), and 15 % chicken embryo extract (CEE, MP Biomedicals™, USA). hMSC aggregates were produced as described by Giger et al.^[53]^ In short, hMSCs were seeded (500 cells per microwell) in Gri3D^®^ 96 well plates containing 121 PEG microwells (⌀ 400 µm) per well according to the manufacturer’s instructions (SunBioscience, Switzerland) in aggregate medium and cultured for 48 h prior to printing. For printing, aggregates were harvested according to the manufacturers protocol, pelleted (300 x *G*, 5 min) and resuspended in PBS.

### Cell casting and bioprinting

Cell-laden constructs were cast to study cell viability in hydrogel formulations containing 10% w/v GelMA with or without 2.5 % w/v PEGDA (Varying M_n_ of 575-10.000), BIS-TRIS (800 mM) or DCPI XC-577 (0.015 % w/v). PEGDA with M_n_ 3,400 and M_n_ 8,000 was obtained from ThermoFisher Scientific, Inc. (USA). For casting, photoresins were prepared essentially as described above, either in sterile conditions (GelMA) or by sterile filtering solutions (PEGDA, BIS-TRIS, DCPI, PBS, LAP) with 0.2 µm PES-filter (Sartorius). PBS was used to replace UPW and 0.25 % w/v of LAP was added to facilitate crosslinking. hMSCs were added in PBS to a concentration of 1x10^6^ cells ml^-1^ and incorporated by careful pipetting. The cell-laden solution was added to a Teflon mold with a connected channel, to cast discs (diameter = 6 mm, height = 2.6 mm). Gels were crosslinked under a glass slide and UV light (365 nm) for 10 min and cultured in hg-DMEM.

To determine compatibility of the Xolography technology with cell-laden photoresins, sterile hydrogels containing 10 % w/v GelMA (X-Pure® GelMA 160P80 RG, Rousselot BV, the Netherlands), 800 mM BIS-TRIS, 1x PBS and 0.015 % w/v DCPI XC-577 were prepared as described. hMSC aggregates were combined with the hydrogel (24 wells ≈ 2904 aggregates per ml gel), transferred to sterile cuvettes for printing and kept at room temperature in the dark to allow for gelation before printing. Printing speed was set to 0.1 mm min^-1^ and UV intensity to 1.2 mW mm^-^². To reduce light scattering, prints were conducted at the side of the cuvette closest to the projector, and a second set was printed on the opposite side after turning the cuvette. Printed samples (diameter = 6 mm, height = 2 mm) were recovered and post-processed as described above and cultured in aggregate medium.

### Metabolic activity assays and fluorescent imaging

AlamarBlue™ Cell Viability Reagent (Invitrogen, USA) was used to determine relative viability of hMSCs in different hydrogel formulations. Three casted constructs were combined with 700 µl of 10 % v/v Alamar Blue in culture medium for 5 h to allow reduction of resazurin into the fluorescent resorufin. Fluorescence was determined in triplicate at 590/35 nm after excitation with 530/25 nm using a BioTek Synergy HTX Multimode plate reader (Agilent Technologies, Inc., USA) and % of reduction was determined based on sample measurements and a medium control. Viability in printed constructs was confirmed using the LIVE/DEAD™ Viability/Cytotoxicity Kit (Invitrogen™). In short, constructs were washed once with PBS before staining with Calcein AM (5 µM) and Ethidium homodimer (10 µM) in PBS for 1 h at 37 °C. Constructs were transferred to coverslips and imaged using an SP5-X confocal microscope using 10x (HCX PL Apo CS, N.A. 0.4) and 20x (HC PL Fluotar L, N.A 0.4) objectives.

### Statistical analysis

Planning and statistical analysis of the DoE was conducted with Statgraphics Centurion 19, (Statgraphics Technologies, Inc., USA). Analysis was performed on the mean with the goal of maximizing, and a medium sensitivity. Parameter impact was pre-set to 3. Two continuous variables were selected with a central composite design (n = 3, centerpoint in triplicate) for hydrogel samples and a 3-level factorial design for UDMA-samples (n = 3, centerpoint in triplicate). A 2^nd^ order regression model was fit through the data and evaluated with two-way ANOVA and lack of fit test (Table S1). Normal probability distribution of the residuals was confirmed for all models. One experimental outlier was removed from the data set for GelMA/PEGDA hydrogels with 5 min UV exposure. PEGDA hydrogels and UDMA prints showed a light-tailed distribution and are sufficient to proceed with the assumption of normality. The two-sided Mann-Whitney U and the two-sided Kruskal-Wallis test were carried out on data obtained for the Hertzian Young’s modulus on milli-scale with GraphPad Prism Version 10.1.2 (GraphPad Software, USA). Results are referred to as statistically significant with p < 0.05 (*) and statistically highly significant with p < 0.01 (**) throughout the work.

### Illustrations and scientific exploitation

Plots were designed with Origin 2023b (OriginLab Corporation, USA), schematics with PowerPoint Version 2406. Contrast and brightness of images was adjusted with Adobe Photoshop 24.0.0 (Adobe Inc., USA) and chemical structures represented by ChemDraw 22.0.0. (Revvity, Inc., USA). Reproduction of the new (Figure 4 A) and old (Figure 4 E, H, I) logo of Eindhoven University of Technology as well as their schematic pixelation (Figure 4 A) is permitted for this purpose under corresponding copyright conditions and by institutional approval.

## Supporting information

VideoSI_GelMA-PEDGA_Resolution

VideoS2_GyroidShrink

VideoS3_BeamBending

## Data Availability Statement

The data that support the findings of this study are available from the corresponding author upon reasonable request.

## Authorship Contribution Statement

**Lena Stoecker:** Conceptualization, Methodology, Investigation, Formal Analysis, Writing – original draft, Visualization **Gerardo Cedillo-Servin:** Methodology, Investigation, Formal Analysis, Writing – original draft **Niklas F. König:** Investigation, Formal analysis, Resources **Freek de Graaf:** Investigation **Marcela García Jiménez:** Investigation **Sandra Hofmann:** Writing –Reviewing and Editing **Keita Ito:** Writing – Reviewing and Editing **Annelieke S. Wentzel:** Investigation, Writing – Reviewing and Editing **Miguel Castilho:** Conceptualization, Methodology, Supervision, Writing – Reviewing and Editing, Funding acquisition

## Acknowledgements

The authors want to express their gratitude to Adriana Gonçalves for valuable discussions and Yasemin B. Atmaca for her assistance in the screening of hydrogel formulations. Further, Baraa Asfari is acknowledged for developing the scripts for grayscaled print file generation. The authors thank Lucíola Vasconcelos Lima, Marcus Reuter, and Dirk Radzinski for their support in 3D printing, processing, and visualization of test structures and for their valuable advice. M.C., S.H., and K.I. acknowledge the funding from the Gravitation program “Materials Driven Regeneration” (023.003.013) funded by the Netherlands Organization for Scientific Research (024.003.013). M.C. and G.C.S. acknowledges support from the EU Horizon 2020 program (grant 874827).

## References

[1] N. Huebsch, P. Arany, A. Mao, D. Shvartsman, O. Ali, S. Bencherif, J. Rivera-Feliciano, D. Mooney, Nature Mater 2010, 9, 518.

[2] C. Yang, M. Tibbitt, L. Basta, K. Anseth, Nature Mater 2014, 13, 645.

[3] S. Callens, D. Fan, I. van Hengel, M. Minneboo, P. Díaz-Payno, M. Stevens, L. Fratila-Apachitei, A. Zadpoor, Nat Commun 2023, 14, 855.

[4] K. Kilian, B. Bugarija, B. Lahn, M. Mrksich, Proc. Natl. Acad. Sci. U.S.A. 2010, 107, 4372.

[5] K. Jung, N. Corrigan, M. Ciftci, J. Xu, S. E. Seo, C. J. Hawker, C. Boyer, Adv Mater 2020, 32, 1903850.

[6] A. Nakayama, Y. Kumamoto, M. Minoshima, K. Kikuchi, A. Taguchi, K. Fujita, Adv Optical Mater 2022, 10, 2200474.

[7] A. Marino, A. Desii, M. Pellegrino, M. Pellegrini, C. Filippeschi, B. Mazzolai, V. Mattoli, G. Ciofani, ACS Nano 2014, 8, 11869.

[8] M. Shusteff, A. Browar, B. Kelly, J. Henriksson, T. Weisgraber, R. Panas, N. Fang, C. Spadaccini, Sci Adv 2017, 3.

[9] L. Rodríguez-Pombo, X. Xu, A. Seijo-Rabina, J. Ong, C. Alvarez-Lorenzo, C. Rial, D. Nieto, S. Gaisford, A. Basit, A. Goyanes, Additive Manufacturing 2022, 52, 102673.

[10] B. Kelly, I. Bhattacharya, H. Heidari, M. Shusteff, C. Spadaccini, H. Taylor, Science 2019, 262, 1075.

[11] P. N. Bernal, M. Bouwmeester, J. Madrid-Wolff, M. Falandt, S. Florczak, N. G. Rodriguez, Y. Li, G. Größbacher, R.-A. Samsom, M. van Wolferen, L. J. W. van der Laan, P. Delrot, D. Loterie, J. Malda, C. Moser, B. Spee, R. Levato, Adv Mater 2022, 34, 2110054.

[12] M. Falandt, P. Nuñez Bernal, O. Dudaryeva, S. Florczak, G. Größbacher, M. Schweiger, A. Longoni, C. Greant, M. Assunção, O. Nijssen, S. van Vlierberghe, J. Malda, T. Vermonden, R. Levato, Adv Mater Technol 2023, 8, 2300026

[13] B. Wang, E. Engay, P. Stubbe, S. Moghaddam, E. Thormann, K. Almdal, A. Islam, Y. Yang, Nat Commun 2022. 13, 367.

[14] H. Liu, P. Chansoria, P. Delrot, E. Angelidakis, R. Rizzo, D. Rütsche, L. Applegate, D. Loterie, M. Zenobi-Wong, Adv Mater 2022, 34, 2204301.

15. [15] V. Hahn, P. Rietz, F. Hermann, P. Müller, C. Barner-Kowollik, T. Schlöder, W. Wenzel, E. Blasco, M. Wegener, Nat Photon 2022, 16, 784.

[16] L. Hafa, L. Breideband, L. Ramirez Posada, N. Torras, E. Martinez, E. Stelzer, F. Pampaloni, Adv Mater 2024, 36, 2306258.

[17] M. Regehly, Y. Garmeshausen, M. Reuter, N. F. König, E. Israel, D. P. Kelly, C. Chou, K. Koch, B. Asfari, S. Hecht, Nature 2020, 588, 620.

[18] L. Stüwe, M. Geiger, F. Röllgen, T. Heinze, M. Reuter, M. Wesseling, S. Hecht, J. Linkhorst, Adv Mater 2023, 36, 2306716.

[19] K. Huang, G. Franchin, P. Colombo, Small 2024, 2402356.

[20] N. Corrigan, X. Li, J. Zhang, C. Boyer, Adv Mater Technol 2024, 2400162.

[21] K. Yue, G. Trujillo-de Santiago, M. Moisés Alvarez, A Tamayol, N. Annabi, A. Khademhosseini, Biomater 2015, 73, 254.

[22] W. A. Green, Industrial Photoinitiators, CRC Press, Boca Raton, 2010.

[23] C. Valderas, S. Bertolotti, C. M. Previtali, M. V. Encinas, J Polym Sci A Polym Chem 2002, 40, 2888.

[24] S. Sharifi, H. Sharifi, A. Akbari, J. Chodosh, Sci Rep 2021, 11, 23276.

[25] R. Goldberg, N. Kishore, R. Lennen, J Phys Chem Ref Data 2002, 31, 231.

[26] W. Huang, R. Yen, M. McLaurine, G. Bledsoe, J Appl Physiol 1996, 81, 2123.

[27] D. Porelli, M. Abrami, P. Pelizzo, C. Formentin, C. Ratti, G. Turco, M. Grassi, G. Canton, G. Grasso. L. Murena, J Mech Behav Biomed Mater 2022, 125, 104933.

[28] C. Greenwood, J. Clement, A. Dicken, J. Evans, I. Lyburn, R. Martin, K. Rogers, N. Stone, G. Adams, P. Zioupos, Bone Rep 2015, 3, 67.

[29] A. Young, O. White, M. Daniele, Macromol Biosci 2020, 20, 2000183.

[30] J. Zatorski, A. Montalbine, J. Ortiz-Cárdenas, Anal Bioanal Chem 2020, 412, 6211.

[31] K. Anseth, L. Kline, T. Walker, K. Anderson, C. Bowman, Macromolecules 1995, 28, 2491.

[32] S. Shin, B Aghaei-Ghareh-Bolagh, T. Dang, S. Topkaya, X. Gao, S. Yang, S. Jung, J. Oh, M. Dokmeci, X. Tang, A. Khademhosseini, Adv Mater 2013, 25, 6385.

[33] J. Lee, H. Lee, EJ Jin, D. Ryu, G. Kim, npj Regen Med 2023, 8, 18.

[34] K. Mamaghani, S. Nahgib, A. Zahedi, M Mozafari, Mater Today 2018, 5, 15635.

[35] Y. Wang, M. Ma, J. Wang, W. Zhang, W. Lu, Y. Gao, B. Zhang, Y. Guo, Materials 2018, 11, 1345.

[36] C. Guimarães, L. Gasperini, A. Marques, R. Reis, Nat Rev Mater 2020, 5, 351.

[37] M. Hippler, E. Blasco, J. Qu, M. Tanaka, C. Barner-Kowollik, M. Wegener, M. Bastmeyer, Nat Comm 2019, 10, 232.

[38] L. Yue, S. Montgomery, X. Sun, L. Yu, Y. Song, T. Nomura, M. Tanaka, H. Qi, Nat Comm 2023, 14, 1251.

[39] A. Puiggalí, R. Rizzo, A. Bonato, P. Fisch, S. Ponta, D. Weber, M. Zenobi-Wong, Adv Healthcare Mater 2024, 13, 2302179.

[40] D. Han, Z. Lu, S. Chester, H. Lee, Sci Rep 2018, 8, 1963.

[41] Y. Jiang, C. Zhu, X. Ma, D. Fan, Biomater Sci 2024, 12, 2504.

[42] L. Rivera-Tarazona, T. Shukla, K. Singh, A. Gaharwar, Z. Campbell, T. Ware, Adv Funct Mater 2022, 32, 2106843.

[43] M. Davidson, M. Prendergast, E. ban, K. Xu, G. ickel, P. Mensah, A. Dhand, P. Janmey, V. Shenoy, J. Burdick, Sci Adv 2021, 7.

[44] M. Zhang, A. Pal, Z. Zheng, G. Gardi, E. Yildiz, M. Sitti, Nat Mater 2023, 22, 1243.

[45] J. Mazzoccoli, D. Feke, H. Baskaran, P. Pintauro, J Biomed Mater Res A 2010, *93A*, 409.

[46] W. Zhu, H. Cui, B. Boualam, F. Masood, E. Flynn, R. Rao, Z. Zhang, L. Zhang, Nanotechnology 2018, 29, 185101.

[47] D. Loessner, C. Meinert, E. Kaemmerer, L. C. Martine, K Yue, P. A. Levett, T. J. Klein, F. P. W. Melchels, A. Khademhosseini, D. W. Hutmacher, Nat Protoc 2016, 11, 727.

[48] Z. Wang, Z. Tian, F. Menard, K. Kim, Biofabrication 2017, 9, 044101.

[49] Y. Garmshausen, M. Reuter, M. Regehly, D. Radzinski (xolo GmbH), EP4224253B1, 2023.

[50] ASTM International, D695–15, USA, 2015.

[51] J. Schindelin, I. Arganda-Carreras, E. Frise, V. Kaynig, M. Longair, T. Pietzsch, S. Preibisch, C. Rueden, S. Saalfeld, B. Schmid, J.-Y. Tinevez, D. White, V. Hartenstein, K. Eliceiri, P. Tomancak, A. Cardona, Nat Methods 2012, 9, 676.

[52] S. Hofmann, H. Hagenmüller, A. Koch, R. Müller, G. Vunjak-Novakovic, D. Kaplan, H. Merkle, L. Meinel, Biomaterials 2007, 28, 1152.

[53] S. Giger, M. Hofer, M. Miljkovic-Licina, S. Hoehnel, N. Brandenberg, R. Guiet, M. Ehrbar, E. Kleiner, K. Gegenschatz-Schmid, T. Matthes, M. Lutolf, APL Bioeng 2022, 8, 036101.

