## Supplementary Information for "Xolography for Biomedical Applications: Dual-color Light-sheet Printing of Hydrogels with Local Control over Shape and Stiffness"

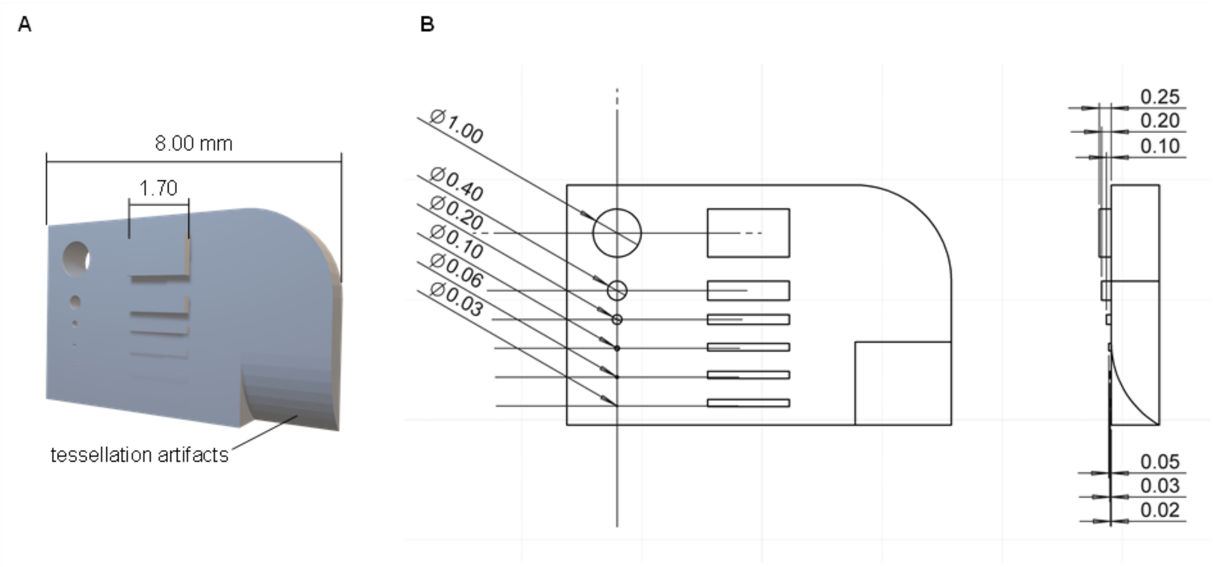


**Figure S1.** The suggested design to assess resolution. (A) 3D view of the sample including tessellation artifacts. (B) Indication of missing dimensions of the CAD model depicted in Figure 3 E, unit is mm.

**Table S1.** Process parameters to determine the processing window including process parameter energy

| Intensity [mW mm$^{-2}$] | Speed [mm min$^{-1}$] | Energy [mJ mm$^{-2}$] |
|---|---|---|
| 0.5 | 0.5 | 3 |
| 0.5 | 1.5 | 1 |
| 0.5 | 2.5 | 0.6 |
| 0.5 | 3.5 | 0.4 |
| 1.5 | 0.5 | 9 |
| 1.5 | 1.5 | 3 |
| 1.5 | 2.5 | 1.8 |
| 1.5 | 3.5 | 1.3 |
| 2.5 | 0.5 | 15 |
| 2.5 | 1.5 | 5 |
| 2.5 | 2.5 | 3 |
| 2.5 | 3.5 | 2.1 |
| 3.5 | 0.5 | 21 |
| 3.5 | 1.5 | 7 |
| 3.5 | 2.5 | 4.2 |
| 3.5 | 3.5 | 3 |

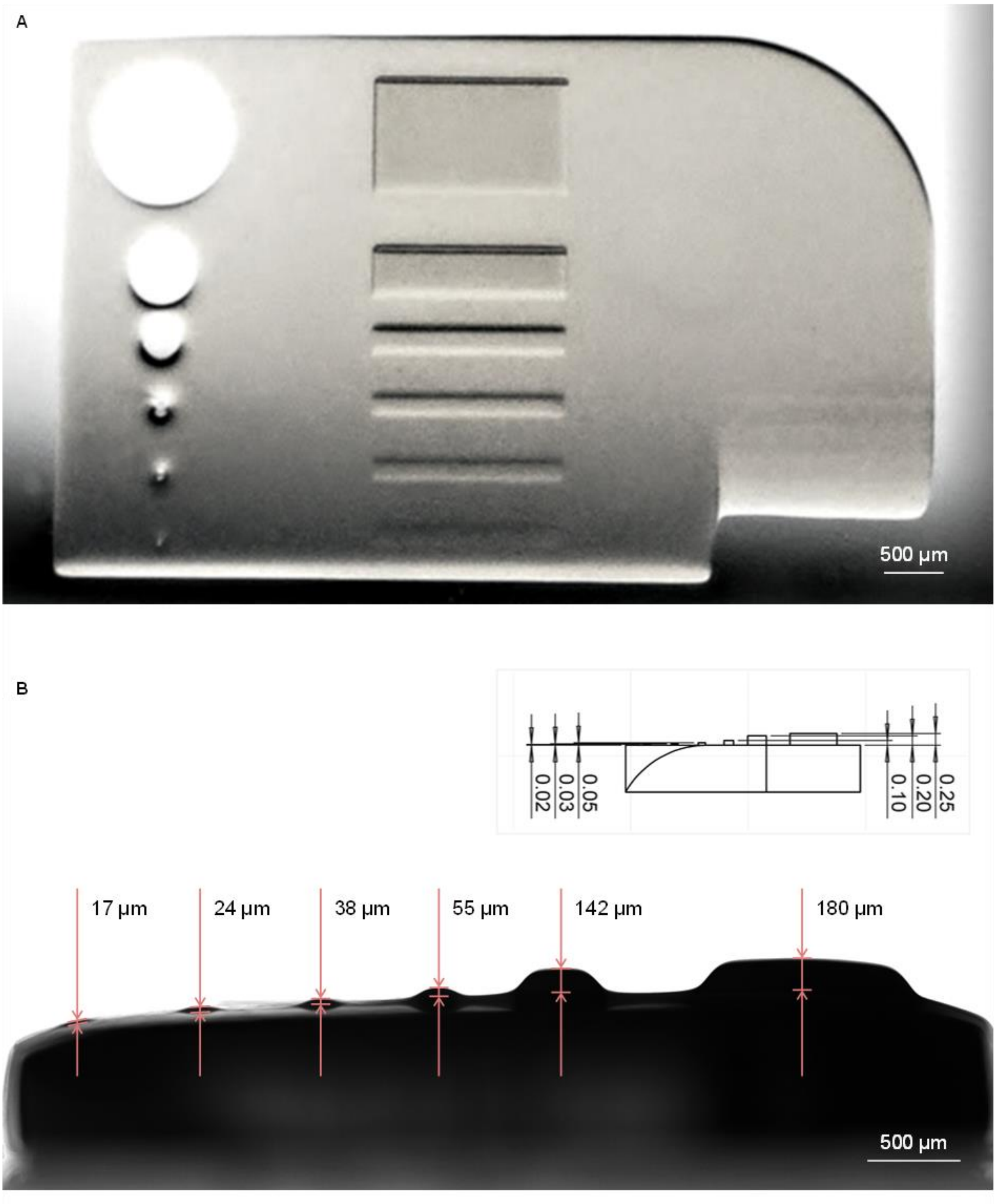


**Figure S2.** Enlarged images of the resolution sample based on the PEGDA-hydrogel. (A) Enlarged version of the image shown in Figure 3 G. (B) Side view showing the formation of positive features with different heights.

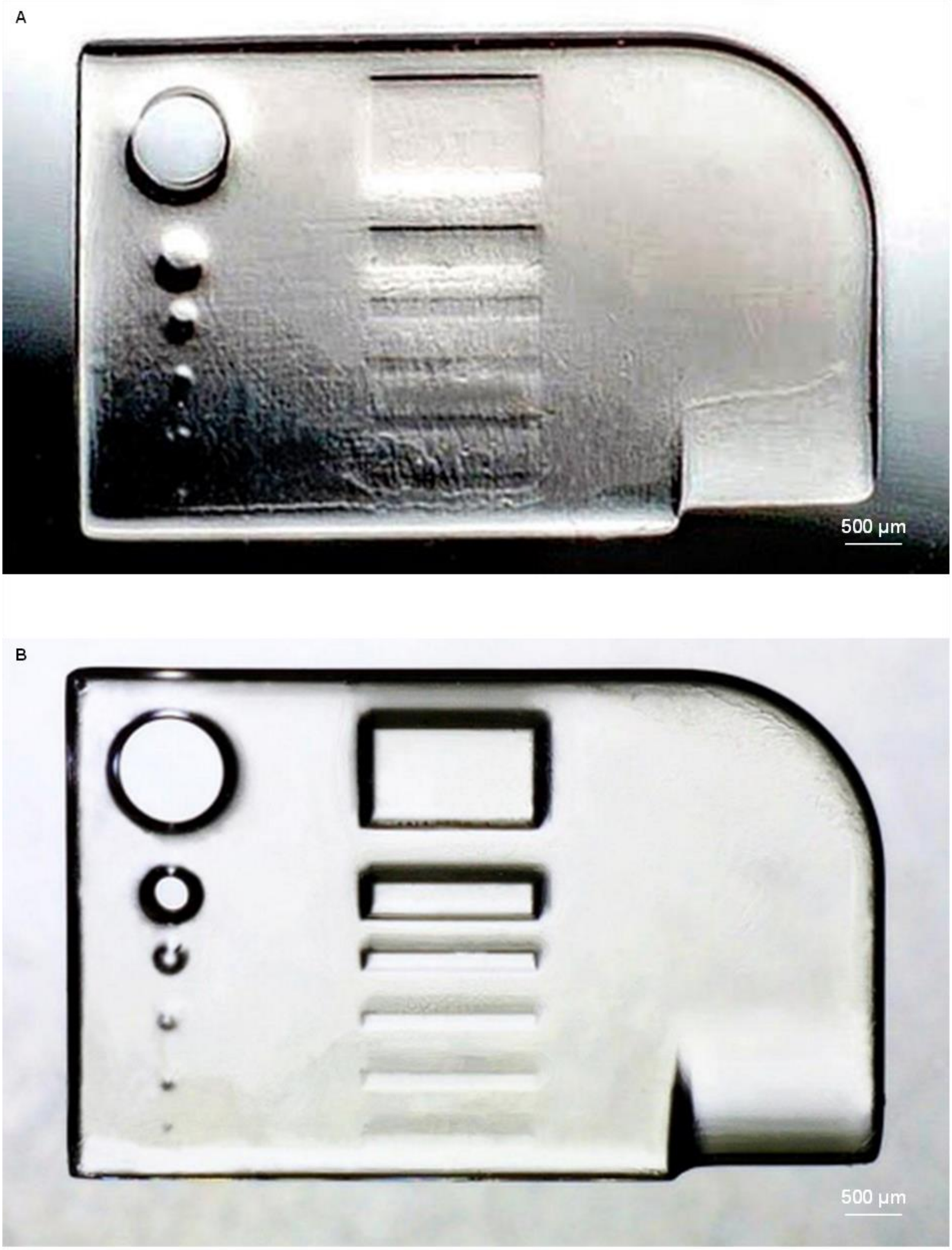


**Figure S3.** Enlarged version of the images of resolution samples based on GelMA/PEGDA and UDMA. (A) GelMA/PEGDA print from Figure 3 F. (B) UDMA print from Figure 3 H.

**Table S2.** Overview of conducted DoE analysis with $p < 0.05$ (*), $p < 0.01$ (**), and $p < 0.001$ (***)

| Photoresin | # | Print direction | Post-processing | $R^2_{adj}$ [%] | Lack-of-fit [ ] |
|---|---|---|---|---|---|
| GelMA/PEGDA | 1 | z | 10 min UV | 63.06 | 0.0699 |
| GelMA/PEGDA | 2 | z | 5 min UV | 81.07 | 0.7218 |
| PEGDA | 3 | z | n/a | 67.17 | 0.8903 |
| UDMA | 4 | z | n/a | 76.21 | 0.3077 |
| UDMA | 5 | z | 5 min UV, 40 min 70 °C | 86.07 | 0.0499* |

**Table S3.** Statistical significance of effects for models containing no lack-of-fit with $p < 0.05$ (*), $p < 0.01$ (**), and $p < 0.001$ (***)

| Photoresin | # | Speed | Intensity | Speed · Intensity | Speed$^2$ | Intensity$^2$ |
|---|---|---|---|---|---|---|
| GelMA/PEGDA | 1 | 0.0009** | 0.1015 | 0.0204* | n/a | n/a |
| GelMA/PEGDA | 2 | < 0.001*** | 0.0341* | n/a | 0.0014** | n/a |
| PEGDA | 3 | < 0.001*** | 0.0004*** | n/a | n/a | n/a |
| UDMA | 4 | < 0.001*** | 0.0015** | n/a | n/a | n/a |

**Table S4.** Regression coefficients for models containing no lack-of-fit

| Photoresin | # | Constant | Speed | Intensity | Speed · Intensity | Speed$^2$ | Intensity$^2$ |
|---|---|---|---|---|---|---|---|
| GelMA/PEGDA | 1 | -0.09 | 0.05 | 0.06 | -0.03 | n/a | n/a |
| GelMA/PEGDA | 2 | 0.01 | -0.01 | 0.0005 | n/a | 0.002 | n/a |
| PEGDA | 3 | 6.64 | -2.03 | 0.32 | n/a | n/a | n/a |
| UDMA | 4 | 6.93 | -20.00 | 2.02 | n/a | n/a | n/a |

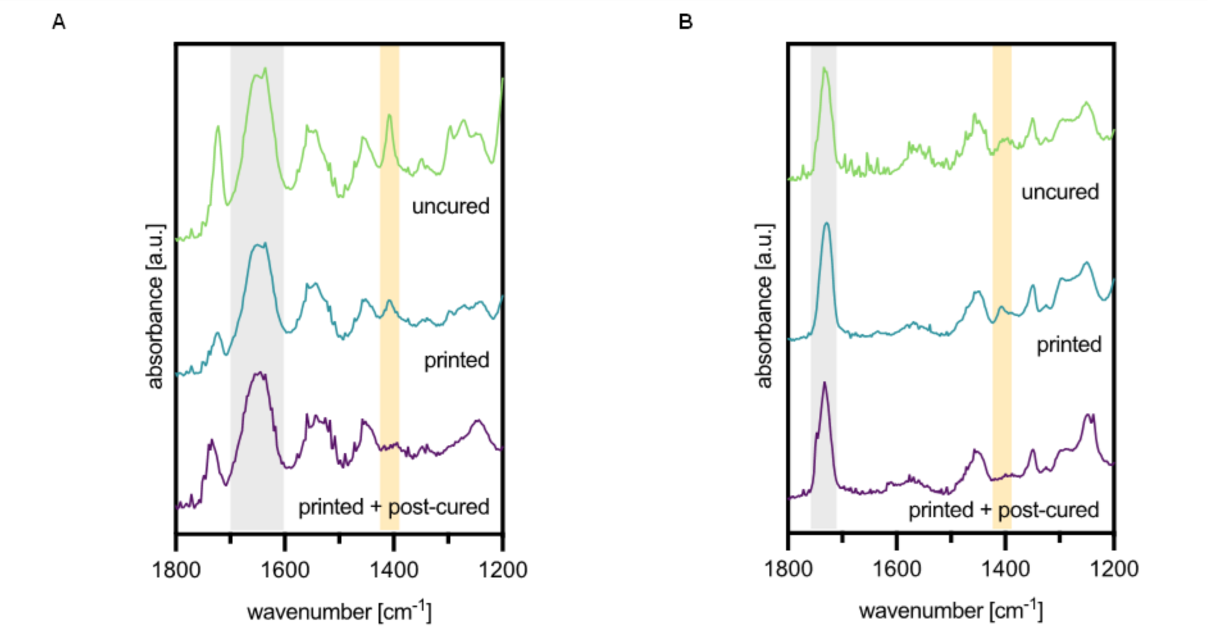


**Figure S4.** FTIR spectra of uncured hydrogel photoresin, printed and post-processed hydrogels. Orange overlay indicates the acrylate C–H scissoring vibration; gray overlay indicates amide II vibration as internal reference. (A) GelMA/PEGDA photoresin. (B) PEGDA photoresin.

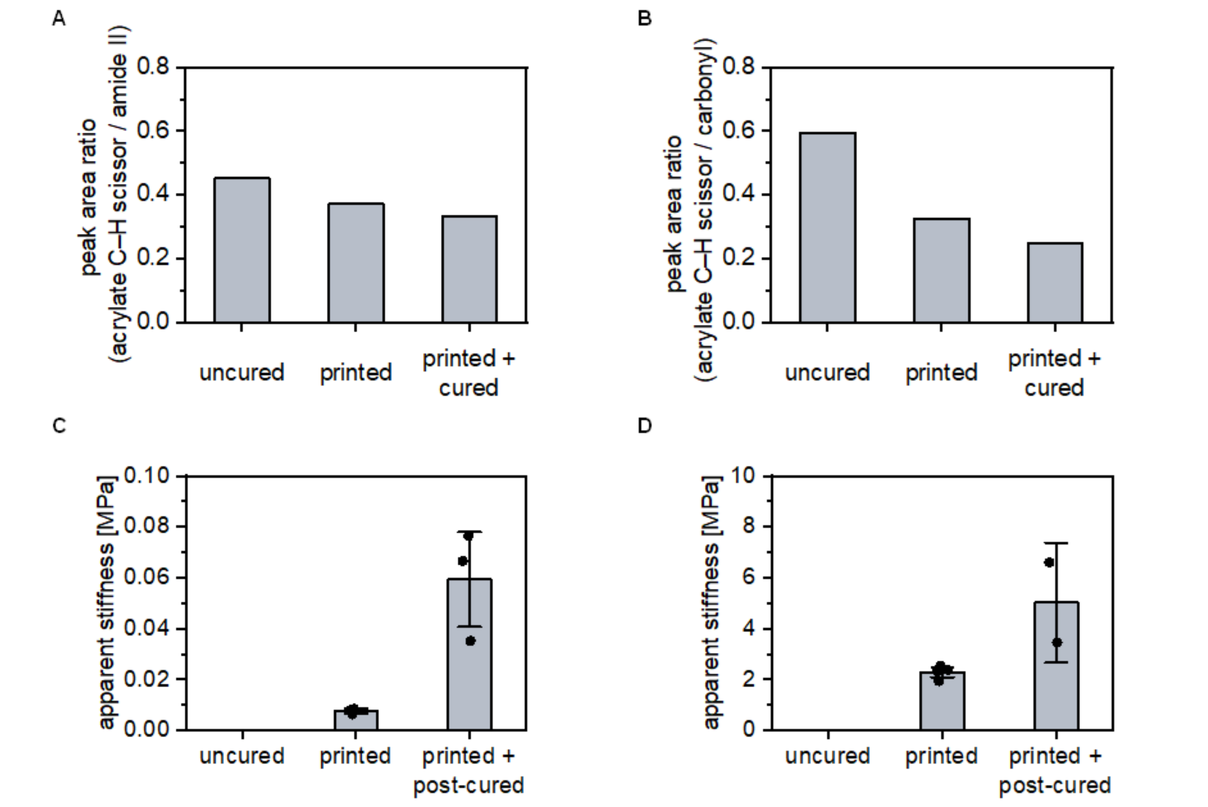


**Figure S5.** (A) Peak area ratio of the uncured photoresin, printed, and post-cured GelMA/PEGDA hydrogels as extracted from FTIR spectra in Figure S2 A. (B) Peak area ratio of the uncured photoresin, printed, and post-cured PEGDA hydrogels as extracted from FTIR spectra in Figure S2 B. (C) Apparent stiffness as obtained from compression tests for printed and post-cured GelMA/PEGDA hydrogels, mean ± SE shown. (D) Apparent stiffness as obtained from compression tests for printed and post-cured PEGDA hydrogels, mean ± SE shown.

**Table S5.** RGB values and the corresponding light intensity as measured at the location of the cuvette during printing

| Short form | RGB code | Intensity [mW cm$^{-2}$] |
|---|---|---|
| RGB 130 | 130, 130, 130 | 119 |
| RGB 190 | 190, 190, 190 | 170 |
| RGB 220 | 220, 220, 220 | 196 |
| RGB 255 | 255, 255, 255 | 242 |

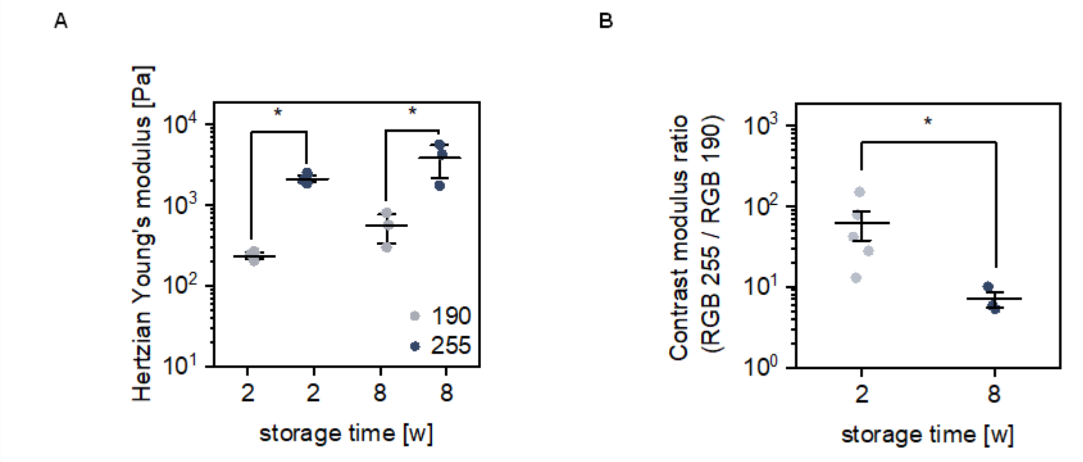


**Figure S6.** Changes in local stiffness and grayscale contrast over time in grayscale-printed PEGDA hydrogels. A) Hertzian Young's modulus after 2 and 8 weeks post-printing, n = 3, two-sided Kruskal-Wallis test, $p \leq 0.0001$, mean ± SE shown. B) The contrast in stiffness defined as the ratio between low and high Hertzian moduli over time, n = 3, two-sided Mann-Whitney U test, $p \leq 0.035$, mean ± SE shown.

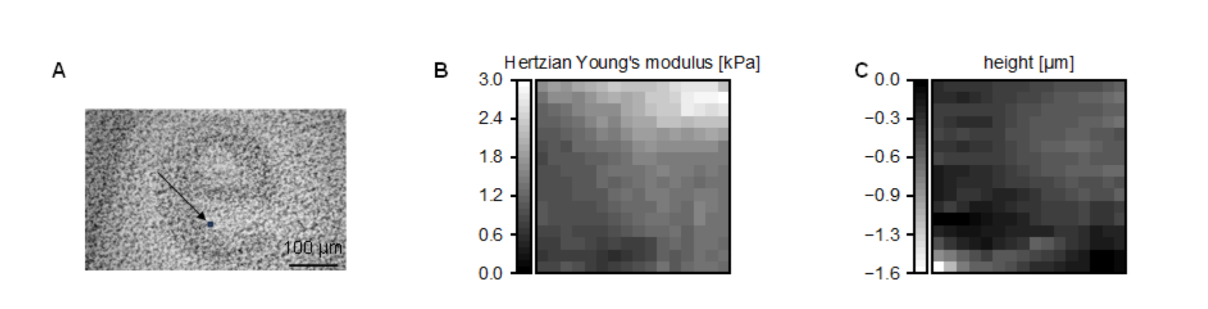


**Figure S7.** Complete data visualization of the stiffness mapping with AFM, A and C are repeated from Figure 4 I and J for clarity. (A) Closeup of the printed text via brightfield, indicating the mapped region (5 µm x 5 µm), n = 1. (B) Hertzian Young's modulus of the region in Figure 4 I and A (RGB = 220). (C) Height map of the region indicated in Figure 4 I and A.

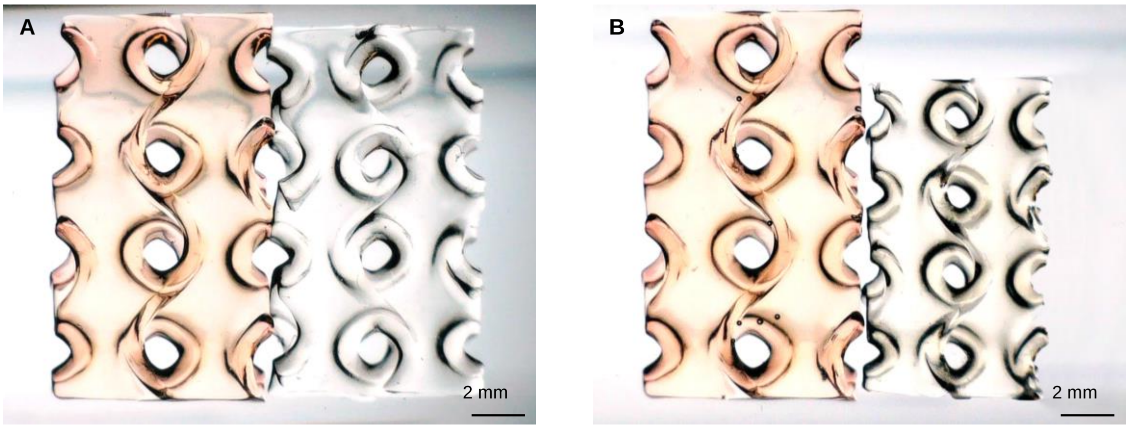


**Figure S8**. Enlarged version of the images in Figure 5 C, showing the gyroid structures printed from GelMA (left gyroid) and NIPAAm/GelMA (right gyroid) photoresins. (A) Gyroids after printing at 20 °C. (B) Gyroids at 50 °C.

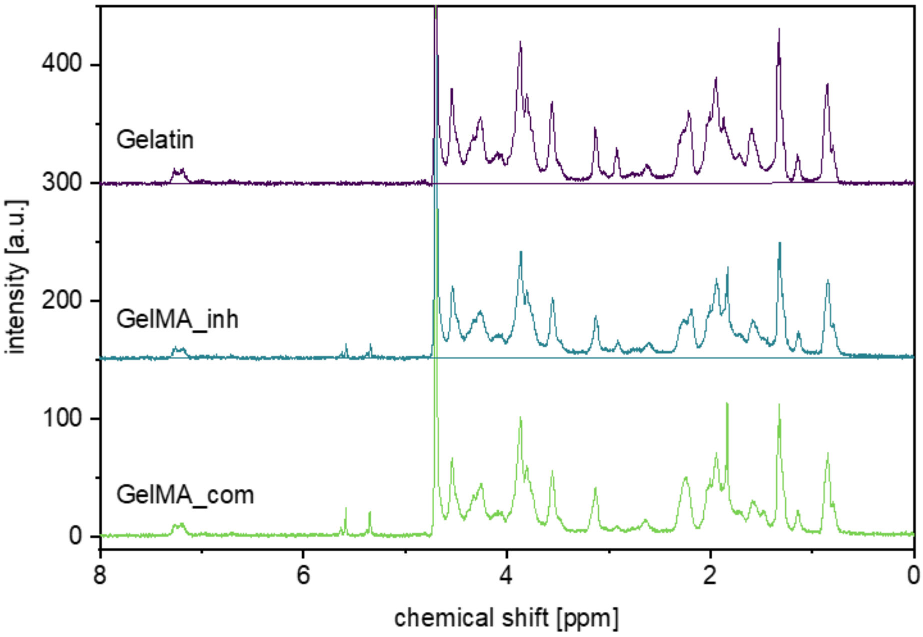


**Figure S90.** [1]H NMR spectrum of gelatin, in-house synthetized GelMA (GelMA_inh), and commercial GelMA (GelMA_com).

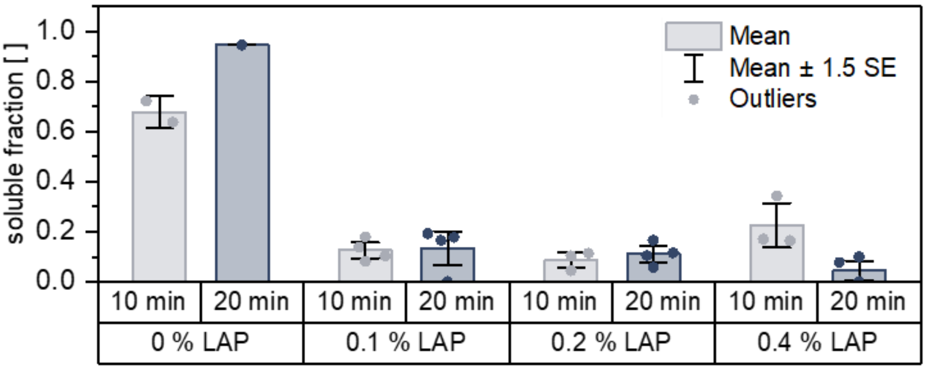


**Figure S10.** Effect of post-processing duration and LAP concentration on sol fraction of printed specimens (n = 3).

**Video S1.** Showing details of the GelMA/PEGDA geometry to assess resolution (visualized in Figure 3 F).

**Video S2.** Shrinking behavior over time at 45 °C for a gyroid structure based on NIPAAm/GelMA hydrogel.

**Video S3.** Partially reversible bending behavior over time of  NIPAAm/GelMA Janus beams printed by grayscaled Xolography, showing anisotropic shape-morphing. Starting from 20 °C, a heating and cooling cycle is shown.
